# Platelets are Protective in Early Abdominal Aortic Aneurysm Formation

**DOI:** 10.1101/2024.10.29.616205

**Authors:** Hannah M. Russell, Anthony R. Spuzzillo, Tyler W. Benson, James Jaworski, Marc Clément, Yacine Boulaftali, Keith Saum, Kelsey A. Conrad, Deborah A. Howatt, James P. Luyendyk, Scott J. Cameron, Bernhard Nieswandt, Steven R. Steinhubl, Alan Daugherty, Wolfgang Bergmeier, Wei Tong, Scott M. Damrauer, Philip S. Tsao, Ziad Mallat, Todd L. Edwards, Nigel Mackman, A. Phillip Owens

## Abstract

**Background:** Abdominal aortic aneurysm (AAA) is a disease associated with the pathophysiologic degradation of the tunica media resulting in aortic dilatation, systemic inflammation, and dysregulated hemostasis. Beyond role its role in initiating primary hemostasis, platelets are a source of ROS, inflammatory cytokines and growth factors necessary for angiogenesis and vascular remodeling. Although platelets contribute to the progression of established aneurysms, their role in the initiation of AAA remains undefined.

**Methods:** Low density lipoprotein receptor deficient (*Ldlr^−/−^*) mice were examined for platelet accumulation in the angiotensin II (AngII) model of AAA utilizing in vivo labeling techniques. Two platelet antagonists (clopidogrel and aspirin), a thrombin inhibitor (dabigatran) or genetic deficiencies (*protease-activated receptor 4*, *P2Y_12_*, *Lnk*) were administered to AngII-infused mice to determine the role of platelets in initiation of AAA. The effect of platelet depletion was examined in multiple mouse strains of AngII-induced AAA and two additional aneurysm models. PheWAS and meta-analysis was analyzed in humans for platelet gene SNPs associated with AAA.

**Results:** We show that platelets are recruited rapidly to the aorta after the initiation of AngII infusion. Genetic deficiency of platelet receptors had no effect on abdominal aortic diameter, but augmented rupture-induced death in littermate versus placebo controls during AngII-induced AAA. Moreover, *Ldlr^−/−^* mice receiving anti-platelet inhibitors or a thrombin inhibitor also had augmented rupture-induced death. Platelet depletion preceding aneurysm formation resulted in pervasive rupture-induced death in several mouse strains and with three different mouse models of AAA.

**Conclusions:** Inhibition of platelet function is detrimental in an early expanding aortic lumen resulting in catastrophic rupture and hemodynamic failure in murine AAA models.

## Introduction

Abdominal aortic aneurysm (AAA) is a degenerative vascular disease characterized by progressive medial and adventitial immune cell infiltration and damage to structural connective tissues giving rise to a permanent localized dilatation of the aortic wall.^1,2^ Expanding asymptomatically until catastrophic rupture, AAA is estimated to afflict more than 1 million people in the US and ~35 million people globally, resulting in >170,000 deaths and 3 million disability-adjusted life years worldwide.^3–5^ Currently, there are no proven pharmaceutical treatments to slow the growth or prevent the rupture of AAA and invasive surgical repair remains the sole interventional option.^6^ While the exact mechanisms of AAA initiation are poorly understood, the earliest events are thought to include an initial vascular lesion which drives localized inflammation, macrophage cell infiltration, and subsequent medial degeneration.^7,8^

Platelets are crucial mediators of hemostasis and are rapid responders during vessel wall injury.^9,10^ Exposed collagen in the subendothelial matrix interacts with platelet glycoprotein VI (GPVI) while collagen-bound von Willebrand factor (vWF) reacts with GP1b, GPV, and GPIX.^11,12,13^ This initial platelet activation is independent of thrombin-mediated agonism, which occurs with tissue factor (TF) dependent activation of the coagulation cascade and culminates with thrombin-mediated cleavage of protease-activated receptors (PARs).^13^ Thrombin-induced platelet activation via PAR1 and PAR4 (humans) and PAR4 (mice) results in the release of adenosine diphosphate (ADP) and thromboxane A_2_ (T_X_A_2_), which activate P2Y_12_ and thromboxane receptors, respectively.^14^ Subsequent dense granule release increases local concentrations of ADP and Ca2+ promoting recruitment and activation of the surrounding platelets to form a platelet plug.^15^ Platelet degranulation also releases matrix metalloproteinases (MMPs) and TGF-β which promote vascular remodeling along with inflammatory cytokines driving leukocyte recruitment.^10^ Platelet activation additionally triggers inside-out signaling leading to conformation changes of integrin receptor αIIbβ3, increasing receptor affinity for fibrinogen and accelerating the formation of a thrombus.^16^

Underpinning the importance of platelets to the pathogenesis of AAA is the formation of a tri-layered intraluminal thrombus (ILT) in approximately 70% of AAAs at the site of expansion.^17,18^ Procoagulant products such as TF, plasmin α2-antiplasmin complexes (PAPs) and d-Dimer are primarily released by the biologically active luminal layer of the thrombus.^19–21^ Platelet activation has been recognized as a contributor to AAA progression by promoting vascular inflammation and propagating growth of the thrombus. Aspirin and P2Y_12_ inhibitors administered to mice with established AAAs attenuate rupture-induced death in growing aneurysms.^22^ Furthermore, the CD41 inhibitor, abciximab, attenuates thrombus size and luminal dilation in a rat xenograft model of AAA.^23^ While platelet inhibitors are associated with decreased aortic dissection and rupture in AAA patients ^24^, several meta-analyses have conversely demonstrated little to no benefit from these drugs.^22,25–27^ Interestingly, additional studies have shown that platelet inhibition can lead to worsening outcomes and rupture in patients with aortic dissection.^28,29^ While platelets likely contribute to the progression of AAAs, the contribution of these cells to the initiation of AAAs is still poorly understood.

Previous work from our group has identified a role for platelet activation in the progression of vascular degeneration and rupture in mouse models of AAA with a confirmatory increased survival in aneurysm patients treated with antiplatelet therapeutics.^22,24,30^ Here, we directly assess the role of platelets in the initiation and early progression of AAA. We identify platelets as an immediate/early responder to the site of aneurysm formation in the angiotensin II (AngII) mouse model of AAA. Genetic (*Par4^−/−^* and *P2y_12_^−/−^*) and pharmacologic (aspirin, clopidogrel, dabigatran etexilate) inhibition of platelets increases the incidence of rupture in a mouse model of AAA. Importantly, platelet depletion results in >80% rupture-induced death in three different AAA models, suggesting platelets are critical for the hemostatic stabilization of AAAs.

## METHODS

### Mice and diet

Male *Ldlr^−/−^*, *apoE^−/−^*, and *C57BL/6J* mice were obtained from The Jackson Laboratory. *Par4^−/−^* mice (Sambrano et al. 2001) were crossed into the *Ldlr^−/−^* background to create *Ldlr^−/−^/Par4^+/+^* and *Ldlr^−/−^/Par4^−/−^*mice. *P2Y_12_^+/+^* and ^−/−^ mice, originally from Portola Pharmaceuticals,^31^ were used as donors in bone marrow transplant experiments. *Lnk*^−/−^ bone marrow was obtained from Dr. Wei Tong, where the mice (originally from Dr. Tony Pawson) are described in previous publications.^32,33^

*ApoE^−/−^* mice were fed a normal laboratory diet throughout experimentation. To induce hypercholesterolemia, *Ldlr^−/−^*mice were fed a diet enriched with saturated milk fat (21% wt/wt) and cholesterol (0.15% wt/wt, Harlan Teklad diet TD.88137 produced by Purina) for 1 week prior to AngII infusion and throughout the duration of infusion.

### Platelet and thrombin inhibition

Platelets were depleted in all mouse models by injecting a rat anti-mouse GP1bα (5 µg/g bodyweight, R300, Emfret Analytics, n = 5) via tail vein, twice weekly for 1 week prior to and throughout AngII infusion. Rat IgG (5 µg/g bodyweight, C301, Emfret Analytics, n = 5) was similarly administered for negative control. Platelet depletion was confirmed by Hemavet analysis (Hemavet 950 LV, Drew Scientific).

For inhibition of thrombin, *Ldlr^−/−^* mice were fed a custom-made HFD containing peanut butter (10 g/kg diet) with or without dabigatran etexilate (10 g/kg diet, purchased from UNC pharmacy: Capsules (150 mg from Boehringer Ingelheim) 1 week prior to and throughout AngII infusion (Dyets Inc., based on Harlan Teklad TD.88137). Dabigatran-induced thrombin inhibition in the plasma was measured with a HEMOCLOT Thrombin Inhibitors kit (Aniara).

For P2Y_12_ platelet inhibition, *Ldlr^−/−^*mice were fed either a custom-made Western diet containing peanut butter flavoring with or without clopidogrel bisulfate (50 mg/kg, purchased from UNC pharmacy: 75 mg Bristol-Meyers Squibb/Sanofi Pharmaceutical tablets) or intraperitoneally injected with clopidogrel bisulfate (30 mg/kg) 1 week prior to and throughout AngII infusion (Dyets Inc., based on Harlan Teklad TD.88137). Clopidogrel bisulfate inhibition of *P2Y_12_* was determined by platelet agonism by ADP ^34^ or tail bleeding time ref. ASA (30 mg/L) was administered to *Ldlr^−/−^* mice via water 1 week prior to and throughout AngII infusion, as described previously ^35^. Effective ASA absorption was determined by plasma levels of thromboxane B2.

### Preparation of macrophage nanoparticles for *in vivo* labeling

Dextran-coated magnetic iron oxide (Fe3O4) nanocrystals were utilized in the in vivo labeling of macrophages, as described previously with modifications (4-6). Aminated (-NH2) detran-coated magnetic iron oxide 20 nm nanocrystals in water were purchased from M K Impex Corp (MKnano, Ontario Canada; product number MKNIOW-DX-020NH2). These nanoparticles were labeled with a DyLight 800 NHS Ester kit (Thermo Scientific, catalog 46421), per the instruction manual. Extraneous dye (270,000 MW) was removed using Vivaspin 6 centrifugal concentrators (Sartorius, 300,000 MWCO). Nanoparticles were further purified by a SuperMag Separator (Ocean NanoTech, incubated overnight per instructions) and resuspended with sterile 1 x Hanks Balanced Salt Solution for in vivo use. Control, unlabeled, nanoparticles underwent the same procedures without addition of DyLight 800 NHS Ester. Labeling was verified by injection into the peritoneum of mice, extraction of peritoneal macrophages 24 hours later, and quantification on an Odyssey LiCor imager with an 800 nm filter.

### Time-course study

*Ldlr^−/−^* mice were fed a Western diet for 1 week prior to and throughout AngII infusion for 2, 4, 7, and 28 days (n = 20 each time point). One week prior to sacrifice, select mice were retroorbitally injected (daily) with an anti-GPIX mouse antibody (emission: 700 nm; 5.0 µg initial loading dose; 2.5 µg total antibody per mouse/day afterward). One day prior to sacrifice, all mice were injected retroorbitally with a solution of dextran-coated nanoparticles (emission: 800 nm; 10 mg final labeled iron oxide/kg body weight), which were created in the above section.

### Aortic imaging and quantification

After sacrifice, aortas were perfused, removed, and immediately cleaned free of all adventitia. Aortas were then placed on an Odyssey Infrared Bioimaging System (LICOR Biosciences) and positioned appropriately. Aortas were then scanned with both the 700 and 800 nm filters on to detect platelets/MMPs or macrophages, respectively. All aortic preparations and scanning parameters remained constant throughout the procedure (Resolution 42 µm; Quality high; Focus offset 1.0 mm; 700 nm channel intensity 5.0; 800 channel intensity 2.0). Aortas were analyzed with the Odyssey software (on-site) or Li-Cor Image Studio Lite version 3.1.4 (remote computer). For quantification of raw platelet and macrophage cells, exact numbers of cells were labeled and analyzed and tissue samples extrapolated. Platelets: non-study *Ldlr^−/−^* mice were injected retro-orbitally with Alexa Fluor 680-labeled GPIX, similar to study animals, for 1 week. Mice were sacrificed and blood was isolated. Platelets were then enumerated using a BD Accuri C6 Flow Cytometer (BD Biosciences). Specific numbers of platelets were then seeded into a 96 well plate (50 µl/well) using a dose curve from 5 x 108 platelets to 0 (Resolution 86 µm; Quality high; Focus offset 3.0 mm; 700 nm channel intensity 7.0). This was performed 5 times and a standard curve was created. Macrophages: non-study *Ldlr*−/− mice were injected intra-peritoneally with DyLight 800-labeled dextran-coated nanoparticles. Twenty-four hours later, macrophages were obtained via peritoneal lavage and enumerated using a hemocytometer. Specific numbers of macrophages were then seeded into a 96 well plate (50 µl/well) using a dose curve from 1.0 x 107 macrophages to 0 (Resolution 86 µm; Quality high; Focus offset 3.0 mm; 800 nm channel intensity 6.0). This was performed 5 times and a standard curve was created. For extrapolation to tissues, abdominal aortas, from the celiac artery to the right renal artery, were excised and cleaned free from adventitia. Aortic segments were then placed in separate solutions of type II porcine pancreatic elastase (250 µg/mL, Sigma) and type I collagen (1 µg/mL, Worthington) for 2 hours at 37°C. All contents were then passed through a 40 µ nylon cell strainer (BD Biosciences) and quantified.

### AngII induction of AAA – Osmotic minipump implantation

At 8 to 12 weeks of age, male mice were implanted with Alzet osmotic minipumps (Model 1004 or 2004, Durect corporation) subcutaneously into the right flank. Infusion of AngII (1,000 ng/kg/min; Bachem) continued for 28 days, as previously described.^36^

### Bone marrow transplantation

This procedure was performed as described previously ^37^. Irradiated mice were re-populated with bone marrow (2 x 10^6^ cells per animal) harvested from *Par4^+/+^* (n = 17), *Par4^−/−^* (n = 16), *P2Y_12_^+/+^* (n = 16), or *P2Y_12_^−/−^* donor mice via retro-orbital injection. Mice were allowed to recover for 4 weeks and were then fed diet and infused with AngII, as referenced in the previous section.

### Deoxycorticosterone acetate (DOCA) salt induction of AAAs

Ten-month old male *C57BL/6J* mice (The Jackson Laboratory) were implanted with DOCA pellets (50 mg, 21-day release; M-121, Innovative Research of America) and concomitantly given salt (0.9% NaCl plus 0.2% KCl), as described previously.^38–40^ Mice were sacrificed at day 28, similar to other mouse experiments.

### Topical elastase model of AAA

C57BL/6J male mice (6 – 8 weeks old, Charles River UK), had porcine pancreatic elastase (Sigma, Type I, ≥4.0 units/mg protein) applied topically to their infrarenal aorta and injected 3 times/week I.P. with 250 μg of the mouse anti-mouse Tgfβ (BioXcell, clone 1D11.16.8) starting on the day of surgery, as described previously.^41^

### Blood pressure measurements

Systolic blood pressure (SBP) was measured on conscious mice using a Coda 8 (Kent Scientific Corporation) tail-cuff system, as described previously ^42^.

### Aortic tissue and plasma collection

Twenty-eight days after pump implantation, mice were terminated, and blood was drawn from the inferior vena cava and plasma processed as described previously ^43,44^. Aortas were perfused with saline, extracted, and placed into formalin (10% wt/vol) until dissection.

### Measurements of abdominal aortic diameters

Abdominal aortas were visualized with high-frequency ultrasound (Vevo 2100, VisualSonics, Toronto, ON, Canada) on day 0 and 27, as described previously.^45,46^ Luminal diameters were measured on images with the maximal dilation. No difference was found between ultrasound measurements and ex-vivo diameter measurements (P = 0.97). An aneurysm is represented by an increase in diameter of 50% or greater from baseline. The mean of suprarenal aortas at baseline is ~0.80 mm, where an aneurysm is represented as a measurement ≥1.20 mm. Represented diameters in publication are from in vivo ultrasound measurements.

### Plasma lipid analysis

Mouse plasma lipids concentrations were analyzed with the following commercially kits: total plasma cholesterol (Total Cholesterol E), triglycerides (L-Type TG M), and HDL-C (L-Type HDL-C) from Wako Chemicals (Richmond, VA). LDL-C was calculated using the Friedewald equation. VLDL-C was then calculated by subtracting HDL-C and LDL-C from total plasma cholesterol.

### Research statistics and data representation

All bar and line graphs were created with Sigma Plot v.15 (SPSS, Chicago, IL). All statistical analysis was performed using SigmaStat, now incorporated into Sigma Plot v.15. Data are represented as mean ± SEM. For two group comparison of parametric data, a Student’s t-test was performed, while non-parametric data was analyzed with a Mann-Whitney Rank Sum. Statistical significance between multiple groups was assessed by One Way analysis of variance (ANOVA) on Ranks with a Dunn’s post hoc, One Way ANOVA with Holm Sidak Post Hoc, or Two-Way ANOVA with Holm Sidak Post Hoc, when appropriate. Statistical significance among groups performed temporally was assessed by either a One-Way Repeated Measures ANOVA (parametric) or Repeated Measures ANOVA on Ranks (non-parametric), where appropriate. Values of *P* < 0.05 were considered statistically significant.

#### Human Study Information

#### Datasets

The BioVU DNA Repository is a deidentified database of electronic health records (EHR) that are linked to patient DNA samples at Vanderbilt University Medical Center. A detailed description of the database and how it is maintained has been published elsewhere.^47^ BioVU participant DNA samples were genotyped on a custom Illumina Multi-Ethnic Genotyping Array (MEGA-ex; Illumina Inc., San Diego, CA, USA). Quality control included excluding samples or variants with missingness rates above 2%. Samples were also excluded if consent had been revoked, sample was duplicated, or failed sex concordance checks. Imputation was performed on the Michigan Imputation Server (MIS, version 1.2.4)^48^ using Minimac4 and the Haplotype Reference Consortium (HRC) panel v1.1.^49^

The MVP study is a large cohort of fully consented participants who were recruited from the patient populations of 63 Department of Veterans Affairs (VA) medical facilities. These results have been previously reported by Giri et al. Genotype calling and QC were performed centrally and genotypes were phased using EAGLE v210 and imputed from the 1000 Genomes Project phase 3 version 5 reference panel using Minimac3 software. The protocols for quality control and analysis to produce the MVP summary statistics were described previously by Klarin et al.^50^

#### Phenome wide association study analysis

We performed a phenome wide association study (PheWAS) of the gene *SH2B3* in the BioVU dataset non-Hispanic whites (N=60,192) and the non-Hispanic blacks (N=13,686), leveraging the full catalog of ICD-9/10 diagnosis codes. We used logistic regression to separately model each of the PheWAS traits (1652 traits for whites, 1263 for blacks) adjusted for sex, age, and 10 principal components. The principal components were derived from a LD pruned set of markers for the two groups. The analysis was run using the PheWAS package in R (version 3.0.6).

#### Meta analyses

The meta-analysis was performed for Abdominal Aortic aneurysm (AAA) outcomes with the BioVU summary stats for AAA (Phecode 442.11) and the AAA summary stats from the MVP study. The program METAL was used to implement the fixed effects, inverse variance weighted meta-analysis for the non-Hispanic white dataset.

#### S-PrediXcan analysis

Genetically predicted gene expression was evaluated for the PheWAS and meta-analysis with S-PrediXcan (PMID: 29739930). S-PrediXcan is a gene-level approach that estimates the genetically determined component of gene expression in each tissue and tests it for association with the outcome using SNP-level summary statistics. The program version MetaXcan was used due its capability to use summary statistics as input. The analysis was run using summary statistics from GTEX-v7. Genetic predictive models significantly predicted *SH2B3* expression brain frontal cortex and brain nucleus accumbens basal ganglia, and were tested for association with AAA risk.

### Study approval

All mouse studies were performed with the approval of the University of North Carolina at Chapel Hill and University of Cincinnati Institutional Animal Care and Use Committee. All analysis of human data was approved by the Institutional Review Board of Vanderbilt University Medical Center.

## RESULTS

### AngII infusion resulted in a time-dependent increase in platelet accumulation at the site of aneurysm formation

We compared platelet and macrophage accumulation during aneurysm formation in AngII-infused *Ldlr^−/−^* mice fed a high fat/high cholesterol diet (HFD/Western) for 1 week prior to and throughout 28 days of AngII infusion (1,000 ng/kg/min). Accumulation measurements using fluorescent labeling of platelets (red) and macrophages (green) were taken on days 0,1, 2, 3, 5, 7, and 28. Platelets rapidly accumulated in the abdominal aorta of AngII-infused mice reaching a significant amount at day 1 and time-dependently increased up to day 28 (Figure 1). Macrophages were recruited with delayed kinetics relative to platelets with first fluorescent signal detection at day 2 and did not reach a significant threshold until day 3 (Figure 1). Mice receiving control agents (IgG and nanoparticles) labelled with 700 and 800 nm reagents had no recordable fluorescent signal (Supplemental Figure 1). Additionally, plasma samples collected from these mice were analyzed for pro-thrombotic and platelet-associated markers. Infusion of AngII resulted in time-dependent increases in plasma concentrations of PF4, TAT, and D-dimer (Supplemental Figure 2). Each blood biomarker evaluated showed a strong positive association with abdominal aortic growth (Supplemental Figure 2).

**Figure 1:**
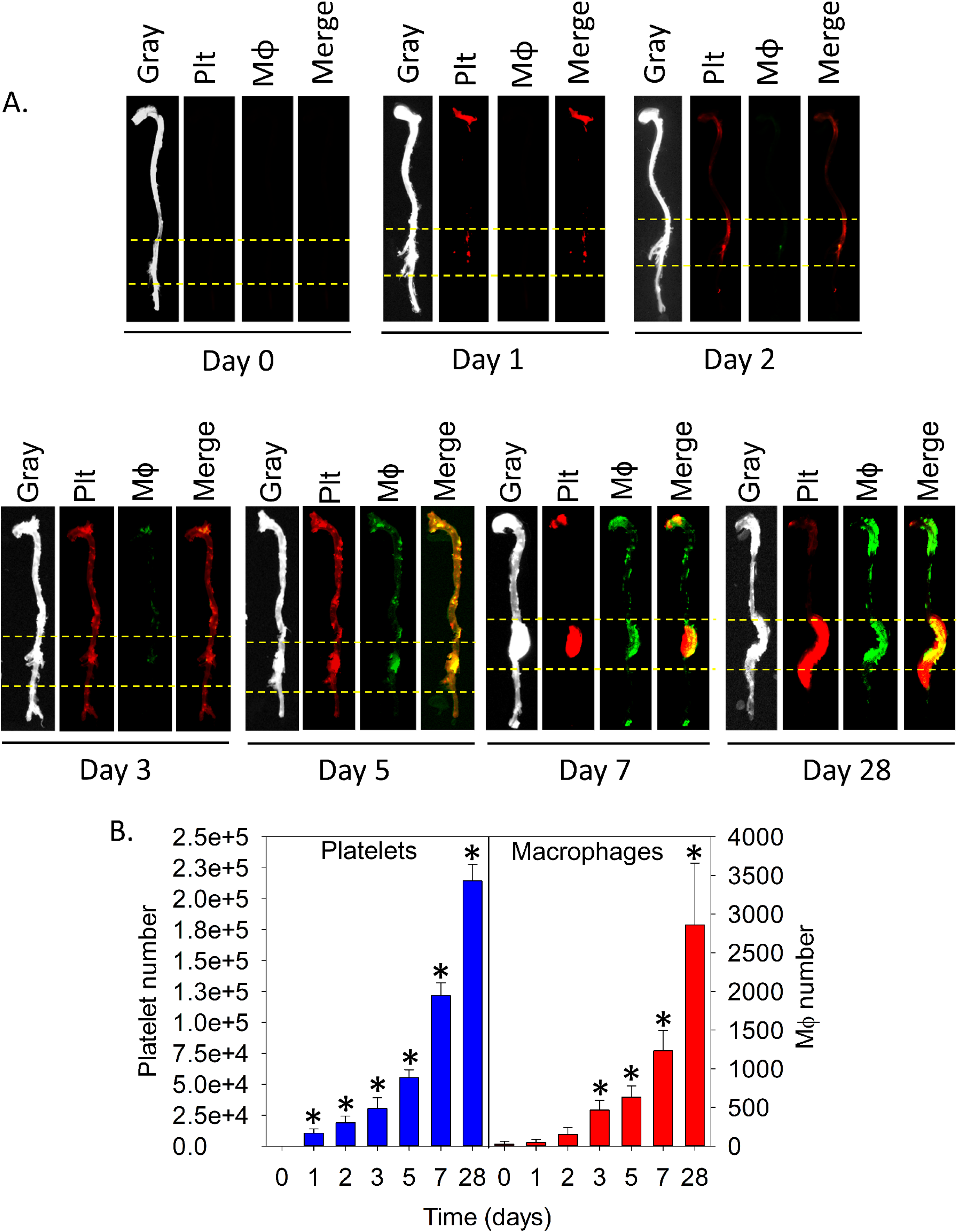
AngII induction of AAA triggered rapid platelet accumulation that preceded macrophage accumulation. Male *Ldlr^−/−^* mice (8 – 10 weeks old) were fed a Western diet for 1 week prior to, and throughout, AngII infusion (1,000 ng/kg/min) for 0, 1, 2, 3, 5, 7, and 28 days (n = 10 – 15 each time-point). Platelets (5 days before sacrifice) and macrophages (24 hours before sacrifice) were labelled with anti-GPIX conjugated 700 nm fluorophore (red) or/and dextran-coated nanoparticles conjugated to DyLight 800 nm fluorophore (green), respectively. (A) Representative platelet, macrophage (MØ), merged, and grayscale images, (B) subsequent quantification. The abdominal aorta within the dotted yellow lines were analyzed for total fluorescent signal in panel B. Histobars represent means ± SEM. (B) *P < 0.05 versus 0 days, repeated measures ANOVA on ranks with Dunn’s post hoc analysis.

### Genetic deficiency of platelet activation receptors augmented AAA rupture

We evaluated the role of PAR4, a key mediator of thrombin-induced platelet activation in mice, in AAA initiation was investigated by utilizing male whole-body deficient *Ldlr^−/−^/Par4^+/+^* and *Ldlr^−/−^/Par4^−/−^*littermates fed a HFD for 1 week prior to, and throughout, AngII infusion for 28 days. While no significant difference in mean aortic diameter was found between *Ldlr^−/−^/Par4^+/+^*and *Ldlr^−/−^/Par4^−/−^* mice, aneurysm incidence and rupture-induced death were significantly increased among *Par4^−/−^* mice (Figure 2A and 2B). To determine if these effects were due to bone marrow (BM)-derived cells (i.e. platelets), *Ldlr^−/−^/Par4^+/+^* and *Ldlr^−/−^/Par4^−/−^*mice were irradiated and repopulated with BM isolated from either *Ldlr^−/−^/Par4^+/+^* or *Ldlr^−/−^/Par4^−/−^* to create 4 chimeric groups. After 4 weeks, successful BM repopulation was verified by isolation and incubation of platelets with PAR4 agonist peptide (P4AP). Platelets isolated from mice receiving *Par4^−/−^* BM were unresponsive to stimulation with P4AP (Supplemental Figure 3A). Similar to whole-body PAR4 deficiency, AngII infusion in mice repopulated with *Par4^−/−^* BM-derived cells resulted in significantly higher rates of rupture-induced death, while mean aortic diameter and aneurysm incidence were similar as compared to control (Figure 2C and 2D). Moreover, administration of the thrombin inhibitor, dabigatran etexilate, (10 g/kg diet) 1 week prior to, and throughout, AngII infusion recapitulated the effects of whole-body PAR4 deficiency in *Par4^+/+^* mice but did not result in any differences in *Par4^−/−^* alone (Figure 2E and 2F).

**Figure 2:**
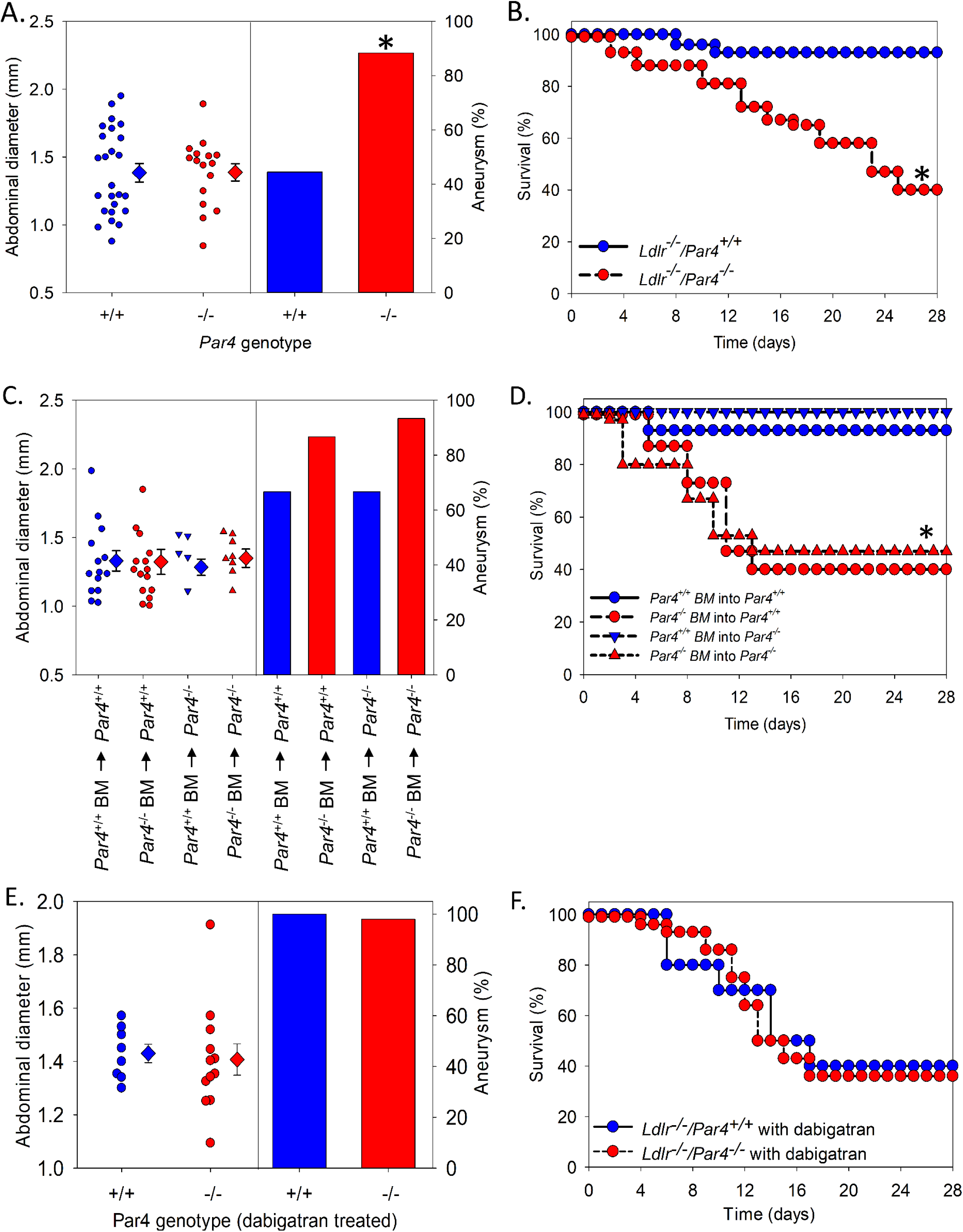
Par4 deficiency augmented AAA rupture via thrombin activation. *Ldlr^−/−^/Par4^+/+^* or *^−/−^* male mice (8 – 10 weeks old) were either irradiated and repopulated with *Ldlr^−/−^/Par4^+/+^* or *^−/−^* bone marrow derived cells (BM; n = 15 each group) and fed a Western diet, or *Ldlr^−/−^/Par4^+/+^* or *^−/−^* male mice (+/+: n = 20; −/−: n = 31) were fed a modified Western diet with dabigatran etexilate (10 g/kg diet) for 1 week prior to, and throughout, AngII infusion (1,000 ng/kg/min) for 28 days. Ultrasonically measured maximal luminal diameters of in vivo suprarenal aortas were measured and AAA incidence determined in PAR4^−/−^ (A), irradiated (C) and dabigatran etexilate fed mice (E). Survival curves were also determined in these groups (B, D, F). Circles represent individual mice, diamonds represent means, and bars represent SEM. Vertical bars represent percent incidence of AAA. *P < 0.001, with both *Par4^−/−^* chimeras versus *Par4^+/+^*chimeras. Data was analyzed with a Fisher’s exact test or a Kaplan Meier estimator.

While PAR1 is the major platelet receptor in humans, it is not expressed on mouse platelets. As such, examination of *Ldlr^−/−^/Par1^+/+^*or *Ldlr^−/−^/Par1^−/−^* littermate mice yielded no difference in aneurysm formation or rupture-induced death (Supplemental Figure 4A and 4B).

Given that platelet activation occurs through multiple independent mechanisms, the role of hematopoietic *P2Y_12_* deficiency in the initiation of AAA was also investigated. Male *Ldlr^−/−^* mice were irradiated, repopulated with BM isolated from either *P2Y^+/+^* or *P2Y_12_^−/−^*mice, and subjected to AngII-induced AAA as detailed above. Mice transplanted with *P2Y_12_^−/−^* BM had markedly reduced platelet activation, as measured by activation of αIIbβ3 upon P4AP or convulxin stimulation compared to mice transplanted *P2Y^+/+^*BM confirming successful engraftment (Supplemental Figure 3C). While there was no difference in abdominal aortic diameter, a non-significant increase in incidence and a significant increase in AAA rupture-induced death was observed in mice receiving *P2Y_12_^−/−^* BM compared to controls (Figure 3). No differences were found between any of the genotypes investigated (PAR4, PAR1, or *P2Y_12_*) regarding body weight, plasma cholesterol, lipoproteins, or systolic blood pressure (Supplemental Table I).

**Figure 3:**
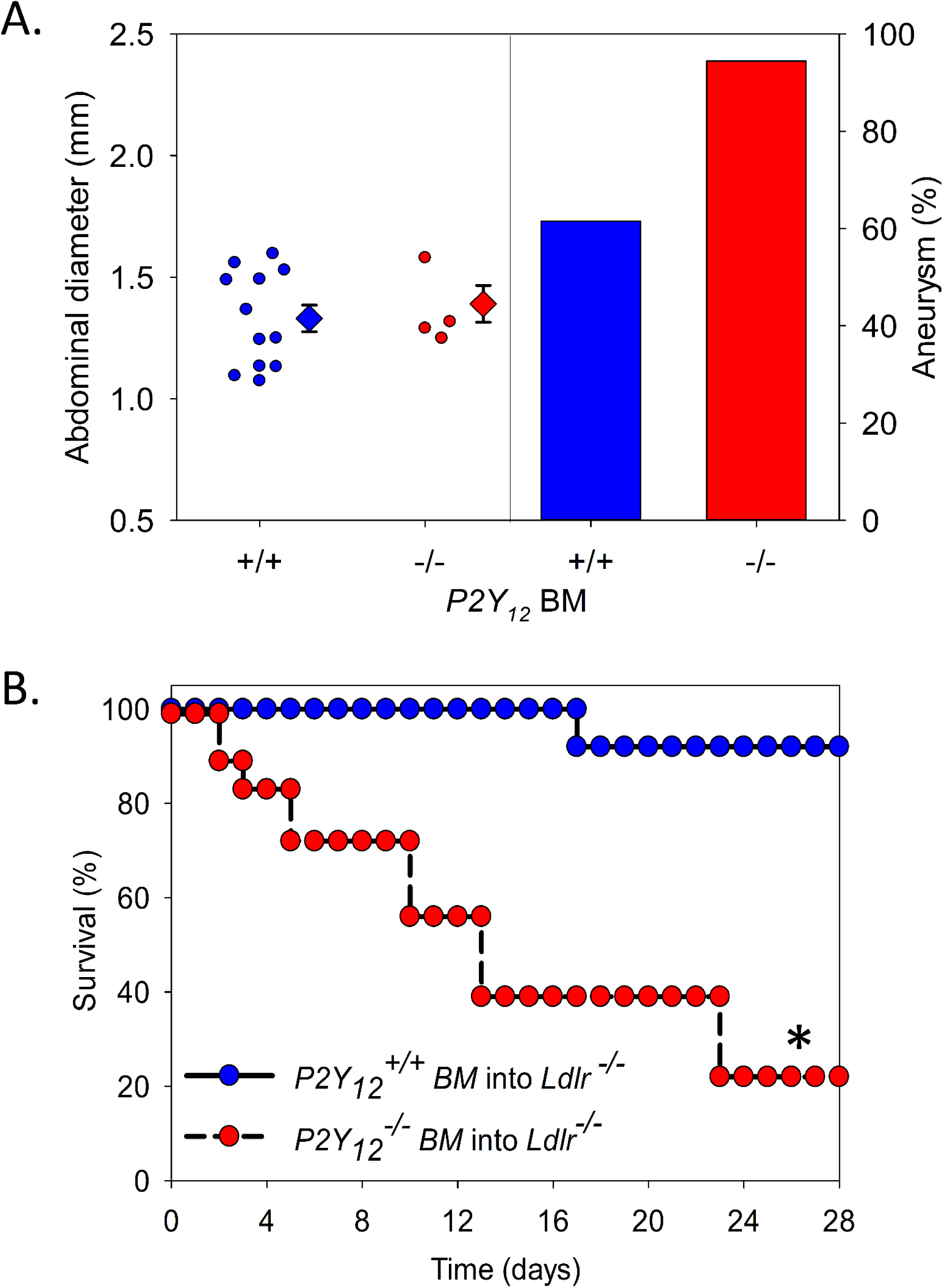
Hematopoietic cell P2Y_12_ deficiency increased AAA rupture. Irradiated *Ldlr^−/−^* mice were repopulated with *P2Y_12_* (+/+: n = 13, −/−: n = 18) bone marrow derived cell (BM) were fed a Western diet for 1 week prior to, and throughout, AngII infusion (1,000 ng/kg/min) for 28 days. (A) Maximal aortic diameters of ex vivo suprarenal aortas were measured and AAA incidence determined in *Ldlr^−/−^ P2Y_12_^+/+^* or *Ldlr^−/−^ P2Y_12_^−/−^* BM. Survival curves were also determined in these groups. Circles represent individual mice, diamonds represent means, and bars represent SEM. Vertical bars represent the percent incidence of AAA. *P < 0.01, *P2Y_12_^−/−^* BM mice versus placebo. Data was analyzed with a Fisher’s exact test or a Kaplan Meier estimator.

### Platelet inhibitors, thrombin inhibition, or platelet deficiency augmented AAA formation and rupture

To determine the role of platelet inhibitors on AngII-induced AAAs, male *Ldlr^−/−^* mice were administered either placebo or ASA (30 mg/L) via drinking water for 1 week prior to and throughout 28 days of AngII infusion. Treatment with ASA completely suppressed thromboxane B2 (T_X_B_2_) secretion into the plasma and reduced integrin activation of isolated platelets when stimulated with arachidonic acid (Supplemental Figure 5A and 5B). In a separate experiment, male *Ldl^−/−^* mice were given clopidogrel bisulfate (oral diet, 50 mg/kg) or vehicle control for 1 week prior to and throughout 28 days of AngII infusion. Platelets isolated from mice fed the clopidogrel bisulfate diet demonstrated almost complete abrogation of ADP-mediated αIIbβ3 activation of ADP stimulated platelets compared to placebo fed mice (Supplemental Figure 5C). Platelet inhibition via ASA or clopidogrel bisulfate had no effect on abdominal aortic diameter and a non-significant increase in AAA incidence (Figure 4A and 4C). Similar to the results of genetic deficiency of platelet receptors, administration of ASA or clopidogrel bisulfate markedly increased rupture-induced death versus placebo controls in the AngII model (Figure 4B and 4D).

**Figure 4:**
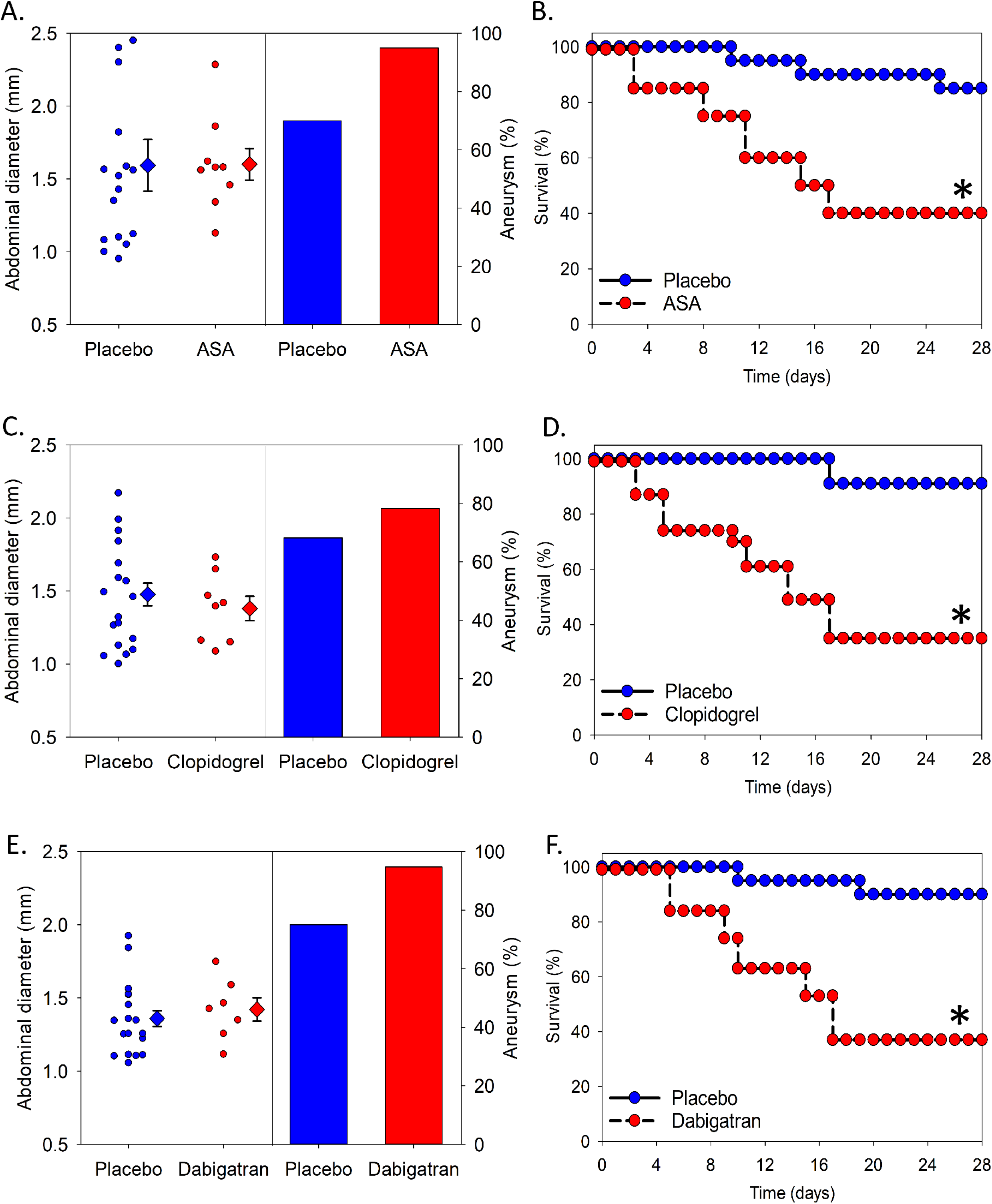
Inhibition of platelet function increased AAA rupture. *Ldlr^−/−^* mice (male; 8 – 10 weeks old) were fed a normal or modified Western diet for 1 week prior to, and throughout, AngII infusion (1,000 ng/kg/min) and administered placebo or acetylsalicylic acid (ASA) via water (n = 20 each group; started with diet), placebo (n = 22) or Plavix (50 mg/kg diet; n = 23) via modified Western diet for 35 days, or placebo (n = 20) or dabigatran etexilate (10 g/kg diet; n = 19) via modified Western diet for 35 days. Ultrasonically measured maximal luminal diameters of in vivo suprarenal aortas were measured and AAA incidence determined in ASA (A), Plavix (C), and dabigatran etexilate administered mice. Survival curves were also determined in these groups (B, D, F). Circles represent individual mice, diamonds represent means, and bars represent SEM. Vertical bars represent percent incidence of AAA. *P < 0.01 drug groups versus controls. Data was analyzed with a Fisher’s exact test or a Kaplan Meier estimator.

Next, the effect of thrombin inhibition on AngII-induced AAA formation was assessed using *Ldlr^−/−^* mice fed HFD containing either placebo or dabigatran etexilate (10 g/kg diet) for 1 week prior to and throughout the experiment. Administration of dabigatran etexilate significantly increased the activated partial thromboplastin time (aPTT; placebo: 0 ng/mL plasma dabigatran; dabigatran etexilate: 289 ± 32 ng/mL plasma dabigatran) indicating effective thrombin inhibition (Supplemental Figure 3B). Similar to direct platelet inhibition, dabigatran etexilate had no effect on abdominal aortic diameter or AAA incidence, but significantly increased the rate of AAA rupture (Figure 4E and 4F). Platelet or thrombin inhibitors had no effect on body weight, plasma cholesterol, lipoproteins, or systolic blood pressure compared to placebo controls (Supplemental Table I).

In contrast to our results, Liu et. al. demonstrated that administration of clopidogrel bisulfate delivered via intraperitoneal injection (IP, 30 mg/kg) to AngII-infused *apoE^−/−^* mice resulted in decreased abdominal aortic diameter and AAA incidence with no change in rupture.^51^ Therefore, we repeated our study with IP administration of clopidogrel bisulfate (30 mg/kg). Similar to our results with oral administration of clopidogrel bisulfate, IP delivered clopidogrel bisulfate had no effect on abdominal aortic diameter or AAA incidence (Supplemental Figure 6A) but increased AAA rupture-induced death (Supplemental Figure 6B). No change in body weight, plasma cholesterol, lipoproteins, or systolic blood pressure was observed (Supplemental Table I). Interestingly, oral administration of clopidogrel bisulfate was more effective at inhibiting platelet function in a tail bleeding assay and integrin activation assay (Supplementary Figure 6C-D).

### Platelet depletion resulted in catastrophic rupture of the abdominal aorta

Since reducing platelet activation through genetic or pharmacologic means augmented rupture-induced death in AngII-infused mice, we next examined the effects of platelet depletion in multiple models of AAA. Platelet depletion with an anti-CD42b antibody (5 µg/g) significantly increased rupture-induced death versus IgG control with no mice surviving past day 12 in hypercholesterolemic *Ldlr^−/−^* mice infused with AngII (Figure 5A). This result was similarly found in another model of hypercholesterolemia (apolipoprotein E – *apoE^−/−^*), and surprisingly, normocholesterolemic *C57BL/6J* mice as well (Figure 5B and 5C). Moreover, platelet depletion in both the deoxycorticosterone acetate (DOCA) salt administered mice (Figure 5D) and elastase exposed mice with concomitant TGFβ inhibition(Figure 5E) recapitulated the severe rupture phenotype.^38^ Taken together, these results suggest that platelets play a protective role in the initiation and stabilization phase of aneurysm formation, regardless of AAA model.

**Figure 5:**
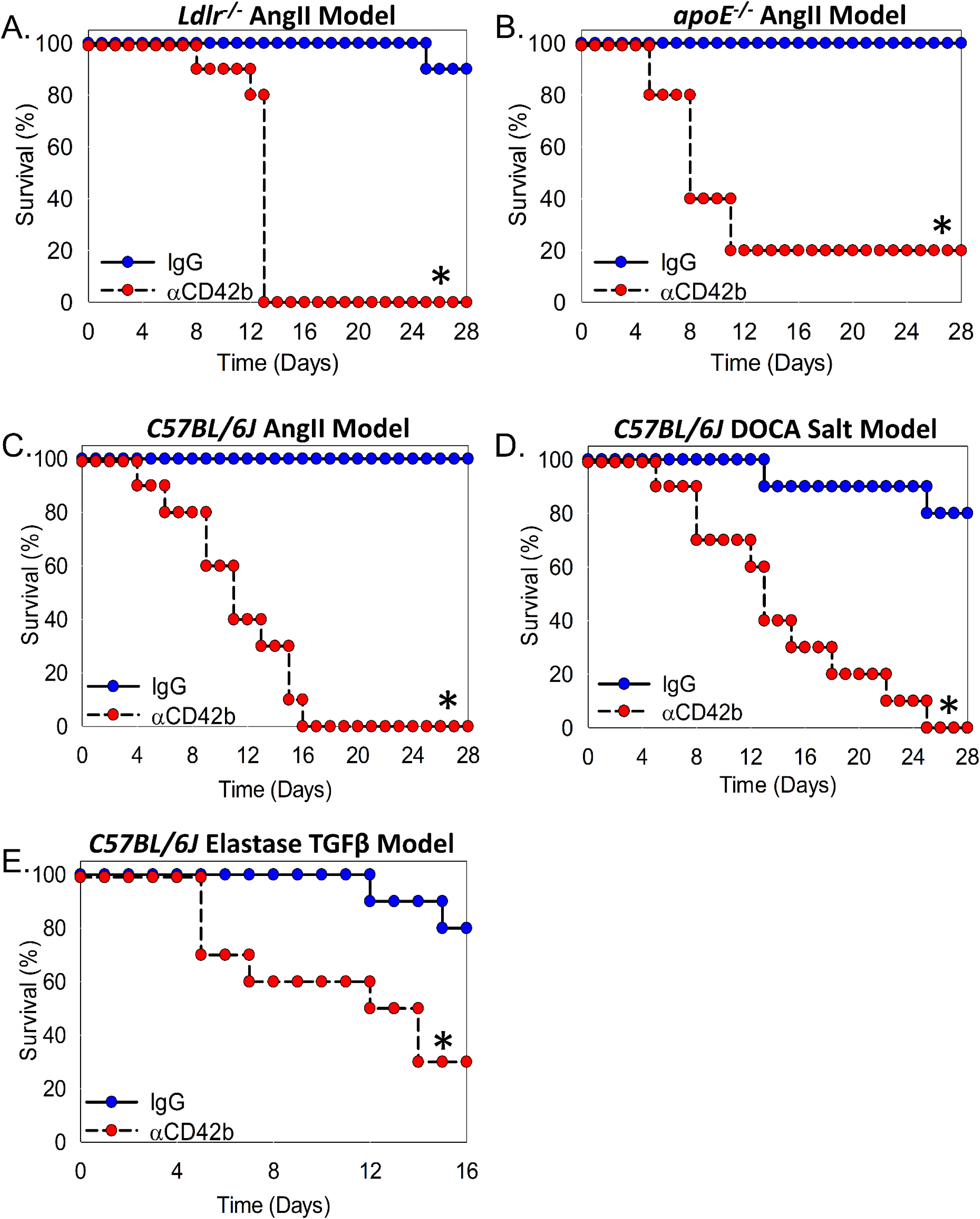
Platelet depletion augmented rupture-induced death in several models of aneurysm formation. Male (A) Male *Ldlr^−/−^* (n = 15 each group), (B) male *apoE^−/−^* (n = 5 each group), and (C) (male, n = 15 each group), (D) (male, n = 10 each group), and (E) male C57BL/6J mice (n = 10 each group; 8 – 12 weeks old; D – 10 months old) were intraperitoneally (I.P.) injected with either non-immune rat IgG or αCD42b platelet depletion antibody (5 µg/g body weight, every 3 days) and (A) fed normal laboratory diet or (B, C) a Western diet for 1 week prior to, and throughout, AngII infusion (1,000 ng/kg/min) for 28 days, (D) implanted with DOCA pellets (50 mg 21-day release) and salt (0.9% NaCl plus 0.2% KCl) in drinking water, (E) had their infrarenal aortas incubated topically with elastase (10 µl 100%) with anti-TGFβ injections (10 mg/kg; 3 times/week, I.P.). Survival curves are represented. *P < 0.05; data was analyzed with a Kaplan Meier estimator.

### Platelet-specific Lnk deficiency was associated with rupture in mice

Lymphocyte adapter protein (LNK), also known as SH2B adapter protein 3, is primarily expressed on the surface of hematopoietic cells, including platelets, and facilitates proper platelet spreading, aggregation, and clot stability by mediating αIIbβ3-dependent outside-in signaling.^16^ GWAS in humans have associated LNK mutations with multiple cardiovascular diseases, including peripheral artery disease (PAD) and coronary artery disease (CAD).^52–55^ Critically, LNK deficiency augments aortic dissection in mice.^56^ To determine the role of LNK deficient platelets in AAA, *Ldlr^−/−^*mice were platelet-depleted with busulfan, and then adoptively transfused with *Lnk^+/+^*, *Lnk^−/−^*, or no platelets every 5 days, with the addition of high fat diet and AngII infusing pumps implanted for 28 days, as described above (Figure 6A). Busulfan administration reduced circulating platelets by 93% in all 3 groups after 16 days (Figure 6B) and a 9% maximal decrease in lymphocytes after 8 days with swift recovery throughout the remainder of the study (Figure 6C). Mice receiving *Lnk^−/−^* platelets or no platelets showed a similar phenotype of catastrophic rupture with complete mortality in both groups by 14 days compared to control mice (Figure 6D).

**Figure 6:**
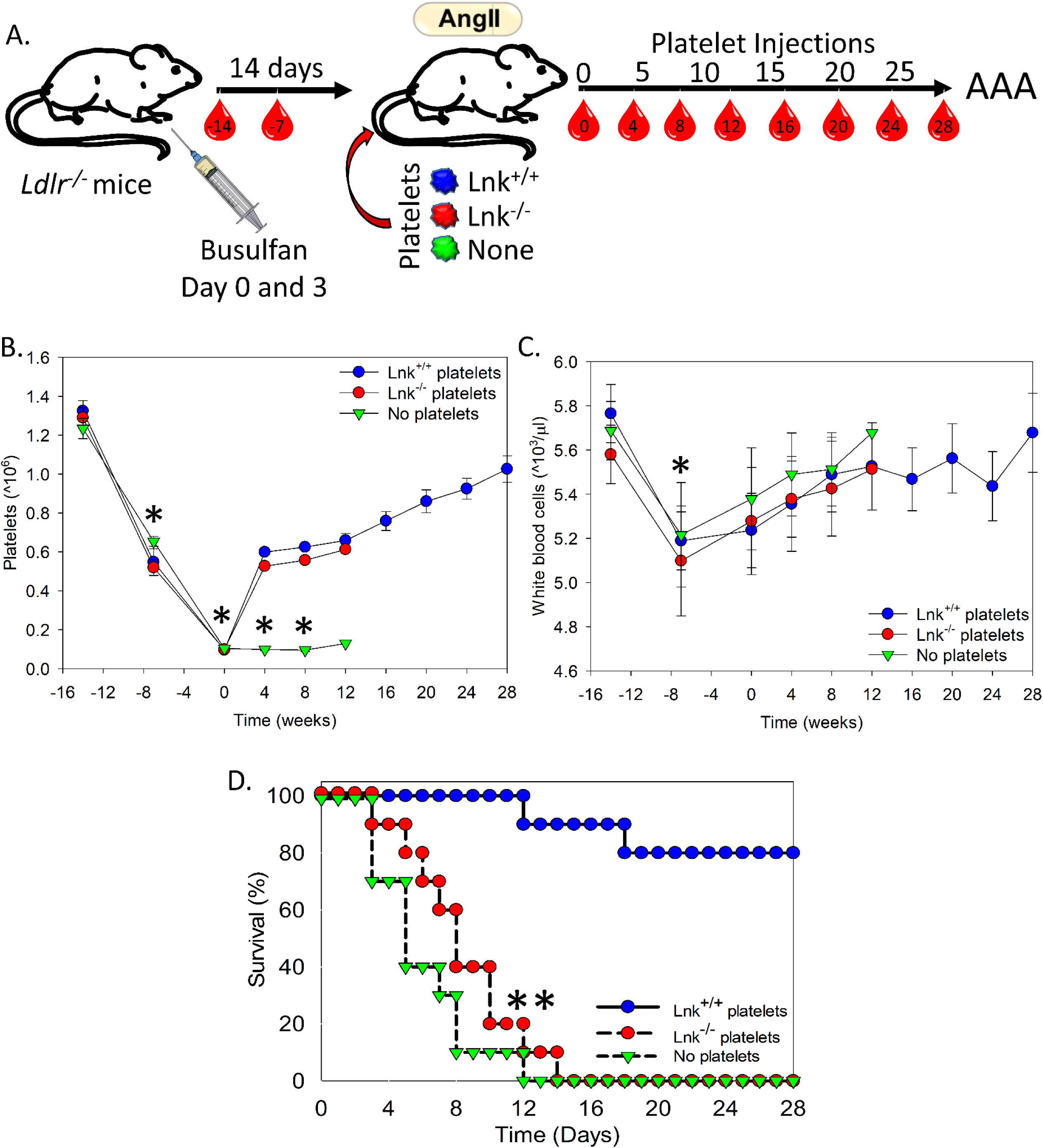
Lnk deficient platelets augmented rupture-induced death. (A) Schematic representation of chronic platelet depletion with busulfan and reconstitution with *Lnk*+/+, *Lnk*−/−, or no platelets. Platelet depleted male *Ldlr−/−* mice were infused with *Lnk*+/+, −/−, or no platelets every 5 days, starting at day 0 with implantation of AngII pumps (1,000 ng/kg/min). Blood was drawn at selected intervals to examine (B) platelets and (C) white blood cells. (D) Representative survival curve during AngII infusion.

### Lnk SNPs associated with augmented aneurysm phenotypes in humans

To determine the susceptibility of *LNK*-related single nucleotide polymorphisms (SNPs) to human aneurysmal disease, we examined both the Vanderbilt BioVU and the Veterans Administration’s Million Veteran Population (MVP) databases (Figure 7A). Meta-analyses were performed for AAA with the Phecode 442.11 and genetically predicted gene expression was further evaluated for the PheWAS and meta-analysis with S-PrediXcan. With the combined databases we found that 8,236 AAA cases and 229,553 controls of both European and African ancestry resulted in 179 unique *SH2B3* SNPs correlated to AAA (Figure 7A). Examination of all patient *SH2B3* PheWAS shows aortic aneurysm and aortic ectasia in the top 1% of relevant ICD-9 Codes (Figure 7B), while the subset of African American PheWAS had aortic ectasia as the fourth-highest significant phenotypic association via ICD-9 code (Figure 7C). Of the 179 unique SH2B3 SNPs associated with AAA, 6 SNPs had significant OR’s >1.870 associated with increased incidence of AAA, where the prevalent cardiovascular disease-associated SH2B3 SNP rs72650673 (OR = 1.847, 95% CI = 1.65 – 2.18, P = 0.0647) nears statistical validity in AAA (Figure 7D).

**Figure 7:**
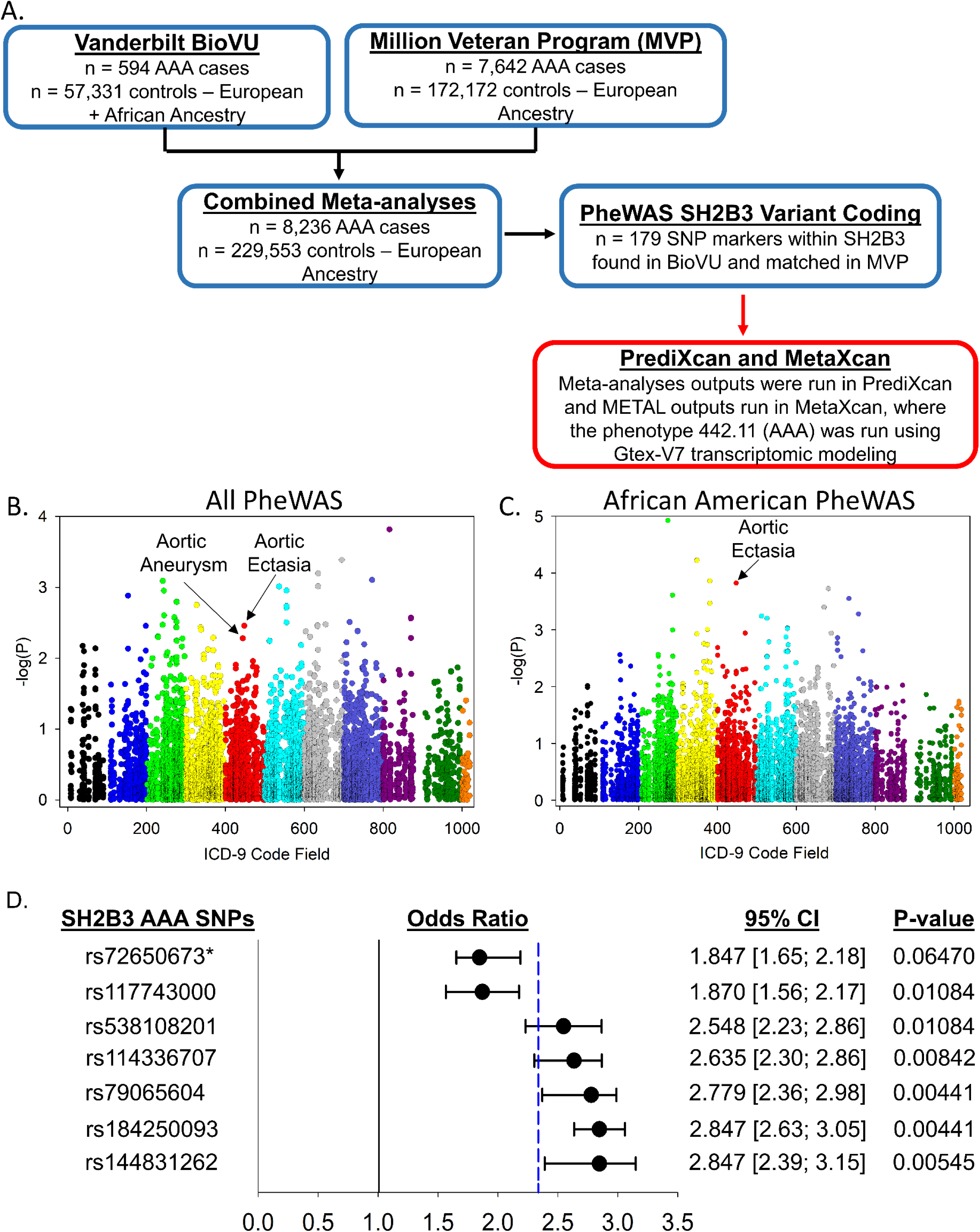
(A) Study design diagram for data sources and analyses for S-PrediXcan PheWAS of SH2B3. (B, C) Plot of −log(p) versus ICD code fields for the PheWAS analysis of genetically predicted SH2B3 expression with risk of diagnoses. (D) Forest plot of associations between SH2B3 SNPs and AAA risk. *P < 0.05 – data analyzed by Repeated Measures ANOVA; **P < 0.05 – data analyzed by Kaplan Meier estimator.

## DISCUSSION

The study of AAA in humans is complicated by asymptomatic onset of the disease posing a serious obstacle to understanding initial conditions and key factors in aneurysm formation. While several rodent studies have demonstrated that intervention with platelet inhibitors results in attenuation of AAA progression of established aneurysms, few studies have examined the role of platelet inhibition prior to the initiation of AAA. This is the first study to demonstrate that platelets are critical for the formation of stable abdominal aortic aneurysms. Herein, we identify several anti-platelet therapeutics (aspirin, clopidogrel, and dabigatran), genetic deficiencies (*Par4^−/−^*, *P2Y_12_^−/−^*, and *Lnk^−/−^*), and platelet depletion in multiple models of aneurysm results in aortic rupture and mortality in mice. To translate these effects into the clinic, we also determine that *LNK* SNPs are associated with aortic aneurysm and ectasia in a large examination of PheWAS datasets. However, collecting data with antiplatelet therapeutics and the initiation of aneurysmal disease in human patients is not possible, as aneurysms are diagnosed only after pronounced dilatation is detected that is likely to be an extended interval after initiation. Regardless, our data, when combined with our prior work demonstrating the protective effects of antiplatelet therapeutics in established aneurysms, suggests platelets have a biphasic role in the pathophysiology of AAA. Platelet accumulation and activation appears to be protective during aneurysm initiation while unchecked propagation in established aneurysms is detrimental. Platelets are considered one of the earliest surveillance cells to detect endothelial injuries as well as microbial pathogens in the vasculature and are likewise among the first to infiltrate the vessel wall in a variety of vascular diseases.^57^ In AAA, we found that PF4, a platelet activation marker, is increased in a time-dependent manner and was significantly correlated with abdominal aortic diameter in AngII-induced aneurysms. While we previously demonstrated plasma PF4 concentrations were significantly increased in AAA patients, no association was found with aneurysm outcomes.^58^ This may be due, in part, to high variability in processing errors with this biomarker.^59^ However, in a more recent analysis, we demonstrate that a more robust and reliable marker of platelet activation, soluble GPVI (sGPVI), is significantly elevated in AAA patients and prognostic of aneurysm diameters and growth rates.^60^ Moreover, it was recently demonstrated that monocyte/macrophage infiltration is necessary for progression to rupture in AngII infused mice.^61^ We speculate that early platelet accumulation may be necessary to prevent monocyte/macrophage induction of rupture by both leading to the chemotaxis of monocyte/macrophages to the site of injury and by providing a robust hemostatic plug and prevention of bleeding diathesis.

In contrast to our results with orally-administered clopidogrel, one study reported that IP-administered clopidogrel reduced aneurysm diameter without affecting rupture in AngII-infused *apoE^−/−^* mice.^51^ As clopidogrel is a pro-drug, it is possible that the drug is not being completely converted to its active form when injected IP.(PMID: 25559342) Indeed, we found that IP injection of clopidogrel had reduced tail bleeding times and activation of platelet integrins when compared to oral administration. However, we still found that IP-administered clopidogrel did increase AAA rupture in *Ldlr^−/−^* mice in a similar manner to orally administered clopidogrel. Furthermore, depletion of platelets in AngII-infused *apoE^−/−^* mice increased rupture, so the discrepancy between the studies is unlikely due to the choice of hypercholesterolemic mouse. While it is easy to point to interlaboratory variability, it is noteworthy that the *apoE^−/−^* platelet depletion experiment was performed at the University of Kentucky, the majority of genetic deficiency models were performed at the University of North Carolina at Chapel Hill, the platelet depletion models were carried out at the University of Cincinnati, and the topical elastase platelet depletion model was performed in Cambridge, England, all with similar results. Together, these results indicate that platelet accumulation and activation are critical in the prevention of rupture at an early stage in multiple mouse models of AAA.

The majority of our experiments utilize the AngII induction of AAAs, which has sufficient parallels to human aneurysm pathology with regard to luminal dilatation, medial elastin breaks, accumulation of atherosclerosis, infiltration of inflammatory cells, proteolytic destruction, intact medial layer, and the penultimate rupture.^7,62^ However, it is also an acute model which results in gross medial rupture of the aortas leading to creation of extraluminal thrombus associated with vascular and adventitial hematoma formation.^7,63^ This hematoma is unlike the ILT typically observed in humans, but rather a thrombus that forms and occludes the expanded lumen and adventitia created by medial rupture and outward pressure exerted by pulsatile blood flow. Yet, several studies have demonstrated that there is corollary blood flow into and out of this extraluminal thrombus, suggesting an exchange with flowing blood similar to the human ILT.^64,65^ We speculate that impaired formation of a stable thrombus allows unchecked expansion and weakening of the false lumen to the point of aortic failure and rupture, demonstrating a critical need for platelet mediated hemostasis at the site of initial aortic dissection. Conversely, after a “stable” aneurysm has formed to resolve the acute vascular injury, continued platelet deposition drives thrombus growth, medial degeneration, progressive inflammation, and overall expansion of the aneurysm, explaining why our lab and others have shown a benefit of platelet inhibition in established aneurysms.^22,30,60^

Additionally, responding platelets are altered in a diseased state. Indeed the platelet transcriptome has significantly differentially expressed genes in AAA patients compared to healthy individuals^30^ and cardiovascular pathology, such as myocardial infarction, increases platelet-derived levels of gelatinase MMP9.^66^ Increased concentrations of MMPs in responding AAA platelets may accelerate proteolytic cleavage of vital structural ECM components potentially worsening established aneurysm, however MMP deficiency in different AAA models has yielded mixed results (PMID: 36795847). Healthy platelets during AAA initiation may be limiting aneurysmal progression and promoting vascular repair through secretion of a variety of factors, such as nitric oxide, tissue inhibitors of matrix metalloproteinases (TIMPs), and CXCL12.^10^

Since it was first associated with celiac disease and type I diabetes,^67^ followed by association with blood pressure and hypertension via a large GWAS,^52^ and more recently cardiovascular disease,^53^ myocardial infarction,^54^ and peripheral artery disease,^55^ missense SNPs in the *SH2B3* gene are highly correlated with augmented disease pathology. Recently, *SH2B3* was associated with aortic dissection utilizing *Lnk^−/−^* mice in the AngII-induced AAAs with human confirmation via PrediXcan association in the BioVU cohort.^56^ Laroumanie and colleagues demonstrated *Lnk* deficiency augmented aortic dissection and rupture via the dysregulation of neutrophils and increased MMP-9 deposition. In addition to *SH2B3* role in lymphoid and myeloid homeostasis, it has also been linked to growth and maturation of megakaryocytes^68,69^ and outside-in signaling of fibrinogen to platelets where *SH2B3* deficiency is associated with poor clot structure and retraction.^16^ Herein, we demonstrate that platelet-specific *Lnk* deficiency also recapitulates the effects of whole body *Lnk* deficiency, and similarly confirms our data suggesting platelet defects results in catastrophic aortic rupture and death in our mouse models. While the majority of our SNPs have been unrecognized in disease pathology, the near significant missense mutation SNP rs72650673 is associated with history of cardiovascular disease (CVD) and CVD risk-factors and altered hematopoietic differentiation, including effects on platelet counts.^53,70^ Similar to the original study by Laroumanie demonstrating SH2B3 gain of function associated with decreased aortic dissection (OR of 0.81 in 229 cases and 16,965 controls), we demonstrate several *SH2B3* loss of function SNPs with ORs 1.8 – 2.8 in 8,236 AAA cases and ~230,000 controls are associated with increased incidence of AAA, with significant PheWAS associations in aortic aneurysm and aortic ectasia.^56^ While their data suggests neutrophils and leukocytes utilizing adoptive transfer into a *Rag1* deficient mouse line, there are links between leukocytes and aggregated platelets which are critical in propagating intravascular thrombosis.^71^ However, this disparity requires further examination.

In summary, this investigation identifies platelets as a critical component of AAA initiation in mice with a link to human pathology. Together, with previous publications, we suggest a dichotomy of platelets in AAA pathology where early activation and deposition of platelets is protective to aortic rupture via either hemostasis preservation or dispersion of forces, while temporal platelet activation and exposure to platelet-derived cytokines and chemoattraction of inflammatory cells results in late-stage pathology and aortic rupture. Future work should be directed toward understanding the mechanism of these dual roles of platelets in aortic pathology.

## Acknowledgements

This work was supported by National Institutes of Health grants 5K99-HL116786, R00-HL116786, R01-HL141404, and R01-HL147171 (APOIII); R01-HL158801-01 (SJC); R35 HL155657 (NM); R35-HL144976 (WB).

## Author Contributions

APO, CJL, SS, VPJ, AE, and STL, CNM conceived and designed the study. CNM, SJC, AM, MG, RB and SKT were responsible for animal care. SJC, SS, DM, SL, MR, AE and RB were responsible for phlebotomy, clinical data management, and regulation. SJC, MG, AA, AM, RB, KK, MTR, DIY, and LW conducted laboratory testing. LW, KK, and DM contributed to the data processing. SJC and CNM supervised all aspects of the study. CNM and SJC contributed to initial data interpretation and wrote the manuscript. All authors contributed to final data interpretation and critical revision of the manuscript, and approved the final version of the manuscript.

