## Supplemental Table 1 and Supplemental Figures 1 - 6 for "Platelets are Protective in Early Abdominal Aortic Aneurysm Formation"

Department of Internal Medicine
[1]Division of Cardiovascular Health & Disease
[2]Pathobiology and Molecular Medicine Graduate Program
[5]Division of Nephrology and Hypertension
University of Cincinnati College of Medicine
Cincinnati, OH 45267-0542, USA

Department of Medicine
[3]Division of Epidemiology, Vanderbilt Genetics Institute, Institute of Medicine and Public Health
Vanderbilt University Medical Center
Nashville, TN 37203

[4]Laboratory for Vascular Translational Science
INSERM
University Paris Diderot
Paris, France

[6]Department of Physiology
Saha Cardiovascular Research Center
University of Kentucky
Lexington, KY 40536, USA

[7]Department of Pathobiology and Diagnostic Investigation
Michigan State University
East Lansing, Michigan 48824, USA

[8]Heart and Vascular Institute, Department of Cardiovascular Medicine, Section of Vascular Medicine
Learner Research Institute
Cleveland Clinic Foundation
Cleveland, OH 44195, USA

[9]Institute of Experimental Biomedicine-Department I,
[10]Rudolf Virchow Center, DFG Research Center for Experimental Biomedicine,
University Hospital Würzburg
97080 Würzburg, Germany.

[11]Weldon School of Biomedical Engineering
Purdue University
206 S. Martin Jischke Drive
West Lafayette, IN 47907, USA

[12]Department of Biochemistry and Biophysics
Department of Medicine
[21]Division of Hematology
UNC Blood Research Center
University of North Carolina at Chapel Hill
Chapel Hill, NC 27599, USA

[13]Division of Hematology, Children's Hospital of Philadelphia
[16]Department of Pediatrics
[14]Department of Surgery
Perelman School of Medicine
University of Pennsylvania, Philadelphia, PA 19104

[15]Corporal Michael J. Crescenz VA Medical Center
Philadelphia, PA 19104, USA

[17]Palo Alto VA Health Care System
Palo Alto, CA 94304, USA

[18]Division of Cardiovascular Medicine, Department of Medicine
Stanford University School of Medicine
Stanford, CA 94305, USA

[19]Division of Cardiovascular Medicine
University of Cambridge
Cambridge, United Kingdom CB2 1TN

[20]Institut National de la Santé et de la Recherche Médicale,
Paris Cardiovascular Research Center
Université Paris-Descartes
Paris, France

*Shared first authorship
**Condensed Title:** Platelets are critical for AAA

**Address for Correspondence:**
A. Phillip Owens III, PhD
University of Cincinnati
231 Albert Sabin Way ML: 0542
Cincinnati, OH 45267-0542


**SUPPLEMENTAL FIGURE LEGENDS:**

**Supplemental Figure 1:** _Control IgG and nanoparticles did not accumulate in AngII infused mice._ Male *Ldlr*$^{-/-}$ mice (8 – 10 weeks old) were fed a Western diet for 1 week prior to and throughout AngII infusion (1,000 ng/kg/min) for 0, 1, 2, 3, 5, 7, and 28 days (n = 10 – 15 each time-point). Inert IgG and nanoparticles labelled with 700 and 800 nm reagents, respectively, were infused as a negative control of background aggregation for the macrophage and platelet accumulation studies. IgG and nanoparticle fluorescent signals were undetectable at each imaging timepoint.

**Supplemental Figure 2:** _AngII induction of AAA increased plasma concentrations of hemostatic proteins correlating with aortic diameter._ Male *Ldlr*$^{-/-}$ mice (8 – 10 weeks old) were fed a Western diet for 1 week prior to, and throughout, AngII infusion (1,000 ng/kg/min) for 0, 1, 2, 3, 5, 7, and 28 days (n = 10 – 15 each time-point). (A) Ultrasonically measured maximal luminal diameters of in vivo suprarenal aortas were measured on days 0, 7, 14, 21 and 28. Citrated plasma was collected from the inferior vena cava and (B) platelet factor 4 (PF4, ng/mL), (D) thrombin-antithrombin (TAT, ng/mL), and (F) D-dimer (ng/mL) were measured by commercially available ELISAs. (C, E, G) Abdominal diameter was significantly correlated with plasma concentrations of hemostatic proteins utilizing the Pearson product-moment correlation coefficient ($4.4 \times 10^{-14} < P < 7.19 \times 10^{-48}$). *P < 0.05 versus 0 days, repeated measures ANOVA on ranks with Dunn's post hoc analysis.

**Supplemental Figure 3:** _Genetic deficiency of platelet receptors impacted platelet activation and clotting._ (A) Platelets isolated from irradiated *Ldlr*$^{-/-}$/*Par4*$^{+/+}$ mice repopulated with *Ldlr*$^{-/-}$/*Par4*$^{-/-}$ bone marrow-derived cells were unresponsive to Par4 agonist peptide (P4AP) via flow cytometry, indicating successful engraftment. (B) Treatment of *Ldlr*$^{-/-}$/*Par4*$^{+/+}$ and *Ldlr*$^{-/-}$/*Par4*$^{-/-}$ mice with dabigatran etexilate increased clotting time in activated partial thromboplastin time (aPPT) tests. *Par4*$^{-/-}$ and dabigatran administration  did not combine to create additional clotting defects. (C) Platelets obtained from irradiated *Ldlr*$^{-/-}$ mice repopulated them with $P2Y_{12}^{+/+}$ or $P2y_{12}^{-/-}$bone marrow were stimulated with P4AP or convulxin. Mice transplanted $P2y_{12}^{-/-}$ bone marrow had significantly reduced integrin activation compared to mice transplanted $P2Y_{12}^{+/+}$ bone marrow derived cells as measured by JON/A via flow cytometry, indicating successful engraftment. Circles represent individual mice, diamonds represent means, and bars represent SEM.

**Supplemental Figure 4:** *Platelet inhibitors reduced markers of platelet activation in AngII infused mice.* Male *Ldlr*$^{-/-}$ mice were given either placebo or a platelet inhibitor 1 week prior to AngII pump implantation and throughout the study. (A) Plasma concentrations of thromboxane B2 ($T_XB_2$) were measured via ELISA in mice given either placebo or 30 mg/L of acetylsalicylic acid (ASA) in drinking water. ASA administration significantly reduced plasma $T_XB_2$ concentrations. (B) Isolated platelets from the two groups were stimulated with arachidonic acid to measure αIIbβ3 integrin activation via flow cytometry. Platelets from mice administered ASA elicited significantly reduced activity when stimulated. (C) Other mice were fed a placebo diet or a diet containing 50 mg/kg of clopidogrel bisulfate. Stimulation of washed platelets with adenosine diphosphate (ADP) showed clopidogrel administered mice had significantly reduced platelet activation compared to placebo fed mice when assessed via flow cytometry. Body weight, plasma cholesterol, lipoproteins, and systolic blood pressure were all unaffected by ASA or clopidogrel platelet inhibition.

**Supplemental Figure 5:** *AngII-induced AAA progression was independent of Par1 receptor in mice.* Male *Ldlr*$^{-/-}$/*Par4*$^{+/+}$ and *Ldlr*$^{-/-}$/*Par4*$^{-/-}$ mice (8 – 10 weeks old) were fed a Western diet for 1 week prior to, and throughout, AngII infusion (1,000 ng/kg/min). (A) No significant difference in aortic diameter or aneurysm incidence was found when comparing *Par4*$^{+/+}$ *and Par4*$^{-/-}$ AngII-infused mice. (B) Rupture-induced death was also unaffected by *Par4* deficiency.

**Supplemental Figure 6:** Male *Ldlr*$^{-/-}$/*Par4*$^{+/+}$ and *Ldlr*$^{-/-}$/*Par4*$^{-/-}$ mice (8 – 10 weeks old) were fed Western diet for 1 week prior to, and throughout, AngII infusion (1,000 ng/kg/min). Mice were given placebo or IP clopidogrel bisulfate injections prior to AngII infusion. (A) Aortic diameter and aneurysm incidence was unaffected by IP clopidogrel, (B) however, there was a significant increase in rupture-induced death in clopidogrel administered mice. Oral clopidogrel administration was a more effective inhibitor of coagulation and platelet activation compared to IP clopidogrel assessed via tail bleed assay and integrin activation assay, respectively (C,D).

**Supplemental Table I: Mice weights, cholesterol levels and systolic blood pressures**

| Mouse Group | Gender | Mouse Number | AAA Model | Treatment | Weight (grams) | TPC (mg/dL) | SBP (mmHg) |
|---|---|---|---|---|---|---|---|
| *$Ldlr^{-/-}$/$Par4^{+/+}$* | Male | 27 | AngII infusion | | 27.49 ± 0.84 | 1358 ± 121.3 | 152.2 ± 8.5 |
| *$Ldlr^{-/-}$/$Par4^{-/-}$* | Male | 31 | AngII infusion | | 28.05 ± 0.68 | 1376 ± 101.2 | 157.4 ± 10.1 |
| *$Par4^{+/+}$ BM* into *$Par4^{+/+}$* | Male | 15 | AngII infusion | | 25.32 ± 0.62 | 1267 ± 102.5 | 148.8 ± 7.8 |
| *$Par4^{-/-}$ BM* into *$Par4^{+/+}$* | Male | 15 | AngII infusion | | 24.85 ± 0.72 | 1251 ± 98.1 | 146.3 ± 9.5 |
| *$Par4^{+/+}$ BM* into *$Par4^{-/-}$* | Male | 15 | AngII infusion | | 25.51 ± 0.49 | 1283 ± 74.5 | 151.1 ± 8.2 |
| *$Par4^{-/-}$ BM* into *$Par4^{-/-}$* | Male | 15 | AngII infusion | | 25.17 ± 0.53 | 1257 ± 87.3 | 146.6 ± 11.2 |
| *$Ldlr^{-/-}$/$Par4^{+/+}$* with dabigatran | Male | 15 | AngII infusion | | 26.34 ± 0.48 | 1684 ± 104.5 | 156.4 ± 6.3 |
| *$Ldlr^{-/-}$/$Par4^{-/-}$* with dabigatran | Male | 19 | AngII infusion | | 26.28 ± 0.37 | 1638 ± 85.6 | 158.7 ± 5.6 |
| *$P2Y_{12}^{+/+}$* BM into *$Ldlr^{-/-}$* | Male | 13 | AngII infusion | | 22.57 ± 0.84 | 1375 ± 75.6 | 141.8 ± 9.6 |
| *$P2Y_{12}^{-/-}$* BM into *$Ldlr^{-/-}$* | Male | 18 | AngII infusion | | 23.01 ± 0.91 | 1370 ± 95.1 | 145.7 ± 6.3 |
| *$Ldlr^{-/-}$* | Male | 20 | AngII infusion | ASA (30 mg/L) | 25.92 ± 0.68 | 1581 ± 96.2 | 154.7 ± 7.6 |
| *$Ldlr^{-/-}$* | Male | 20 | AngII infusion | Placebo | 26.14 ± 0.74 | 1524 ± 113.5 | 158.9 ± 8.2 |
| *$Ldlr^{-/-}$* | Male | 22 | AngII infusion | Clopidogrel Bisulfate (Oral, 50 mg/kg) | 24.82 ± 0.51 | 1684 ± 74.2 | 157.9 ± 9.5 |
| *$Ldlr^{-/-}$* | Male | 23 | AngII infusion | Placebo | 24.74 ± 0.46 | 1653 ± 84.1 | 155.6 ± 12.3 |
| *$Ldlr^{-/-}$* | Male | 19 | AngII infusion | Dabigatran etexilate (10 g/kg diet) | 26.81 ± 0.58 | 1624 ± 52.6 | 164.7 ± 5.6 |
| *$Ldlr^{-/-}$* | Male | 20 | AngII infusion | Placebo | 27.02 ± 0.93 | 1651 ± 62.8 | 159.9 ± 10.5 |
| *$Ldlr^{-/-}$* | Male | 12 | AngII infusion | Clopidogrel bisulfate (IP, 30 mg/kg) | 25.57 ± 0.59 | 1592 ± 75.6 | 157.6 ± 7.3 |
| *$Ldlr^{-/-}$* | Male | 12 | AngII infusion | Placebo | 25.86 ± 0.47 | 1621 ± 105.8 | 156.9 ± 6.1 |
| *$Ldlr^{-/-}$/$Par1^{+/+}$* | Male | 15 | AngII infusion | | 24.25 ± 0.52 | 1785 ± 85.2 | 146.8 ± 7.8 |
| *$Ldlr^{-/-}$/$Par1^{-/-}$* | Male | 15 | AngII infusion | | 24.17 ± 0.67 | 1821 ± 58.5 | 149.5 ± 6.5 |
| *$Ldlr^{-/-}$* | Male | 15 | AngII infusion | anti-CD42b antibody (5 µg/g) | 25.08 ± 0.87 | 1681 ± 69.3 | N/A |
| *$Ldlr^{-/-}$* | Male | 15 | AngII infusion | placebo | 24.86 ± 0.93 | 1721 ± 86.2 | N/A |
| *$apoE^{-/-}$* | Male | 5 | AngII infusion | anti-CD42b antibody (5 µg/g) | 25.27 ± 0.75 | 486 ± 42.6 | N/A |
| *$apoE^{-/-}$* | Male | 5 | AngII infusion | placebo | 25.14 ± 0.68 | 512 ± 35.8 | N/A |

| | | | | | | | |
|---|---|---|---|---|---|---|---|
| *C57BL/6J* | Male | 15 | AngII infusion | anti-CD42b antibody (5 µg/g) | 25.12 ± 0.73 | 158 ± 26.3 | N/A |
| *C57BL/6J* | Male | 15 | AngII infusion | placebo | 24.94 ± 0.48 | 146 ± 18.6 | N/A |
| *C57BL/6J* | Male | 10 | DOCA Salt administered mice | anti-CD42b antibody (5 µg/g) | 34.85 ± 1.43 | 95 ± 8.2 | N/A |
| *C57BL/6J* | Male | 10 | DOCA Salt administered mice | placebo | 35.17 ± 1.86 | 91 ± 7.7 | N/A |
| *C57BL/6J* | Male | 10 | Elastase + TGFβ Inhibition | anti-CD42b antibody (5 µg/g) | 25.74 ± 0.42 | N/A | N/A |
| *C57BL/6J* | Male | 10 | Elastase + TGFβ Inhibition | placebo | 25.62 ± 0.38 | N/A | N/A |
| *Ldlr*$^{-/-}$ | Male | 12 | AngII infusion | *Lnk*$^{+/+}$ platelet transfusion | 24.05 ± 0.75 | 1574 ± 82.3 | N/A |
| *Ldlr*$^{-/-}$ | Male | 12 | AngII infusion | *Lnk*$^{-/-}$ platelet transfusion | 23.84 ± 0.68 | 1526 ± 79.5 | N/A |
| *Ldlr*$^{-/-}$ | Male | 12 | AngII infusion | No platelet transfusion | 23.67 ± 0.54 | 1602 ± 102.3 | N/A |

### Supplemental Figure 1

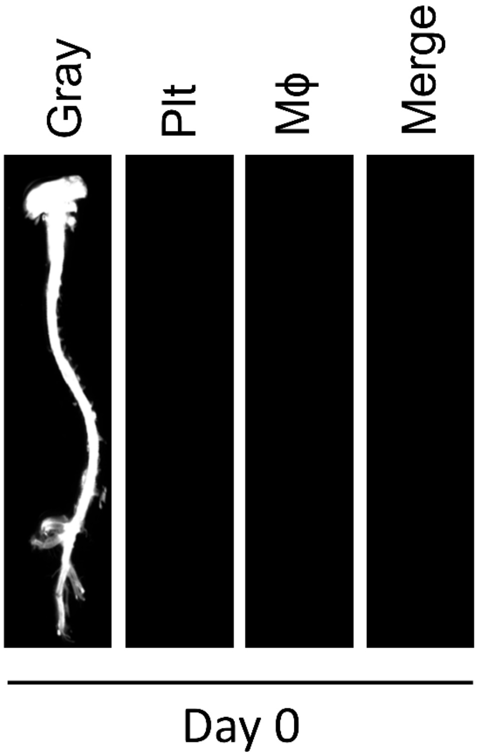

### Supplemental Figure 1 (continued)

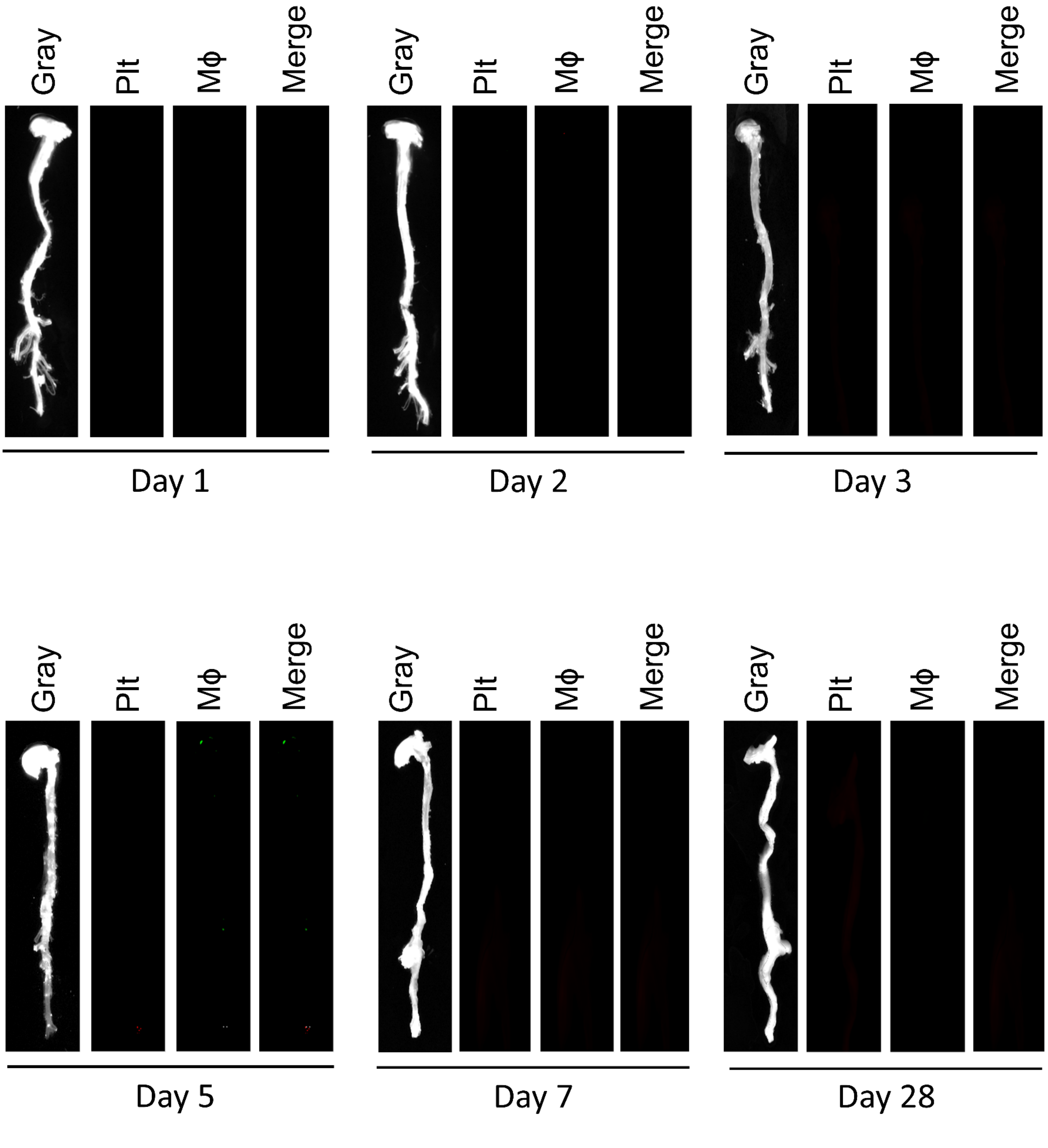

**Supplemental Figure 2**

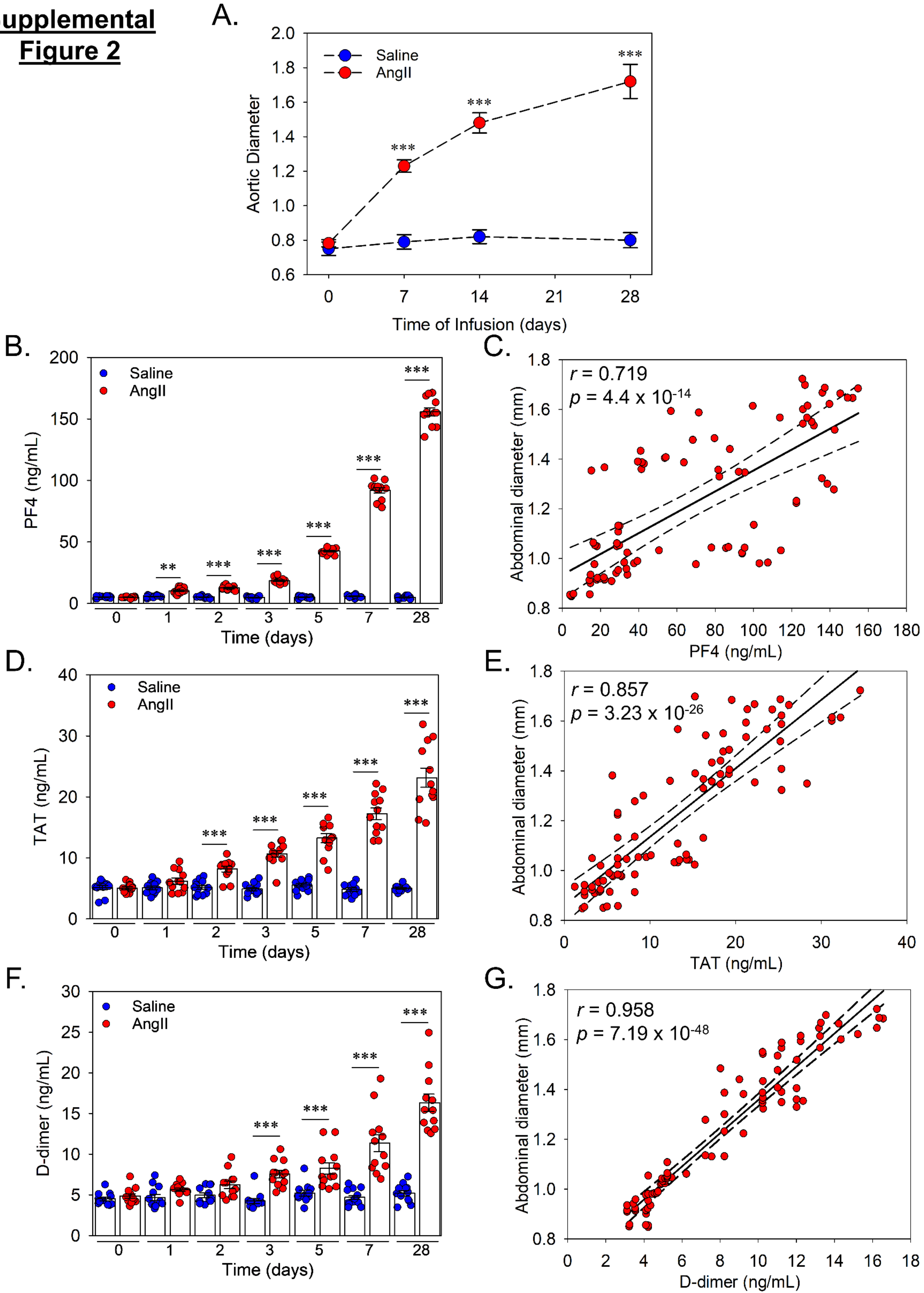

**Supplemental Figure 3**

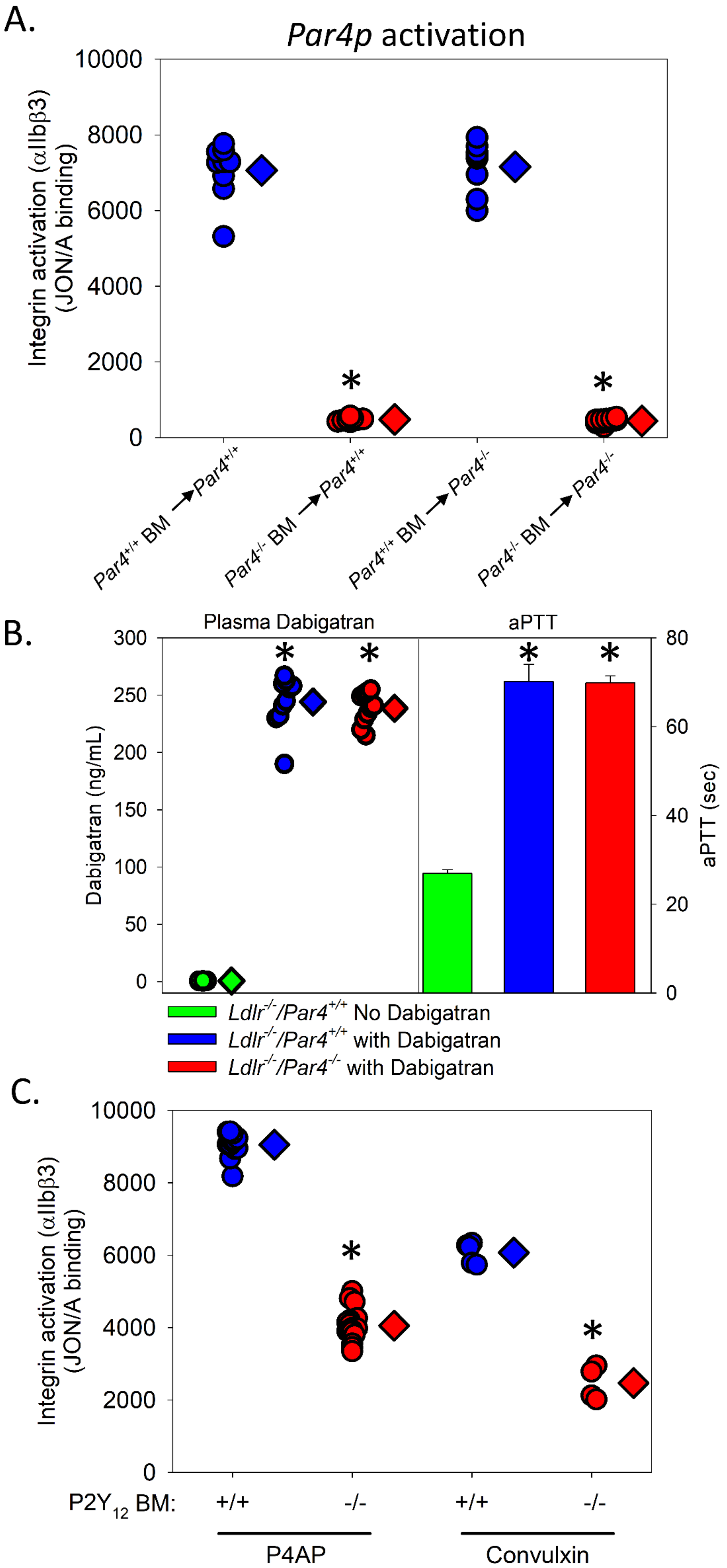

**Supplemental Figure 4**

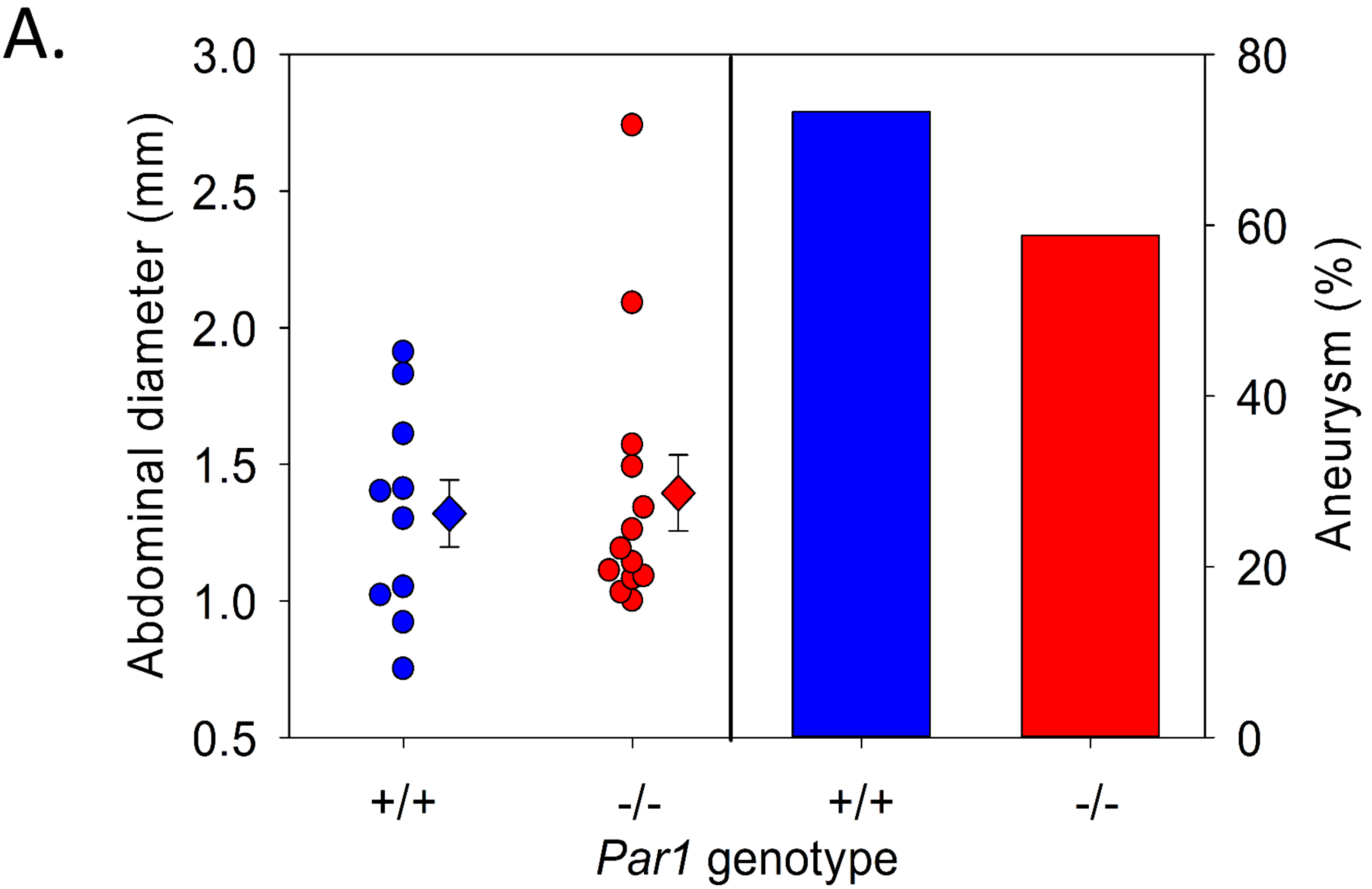


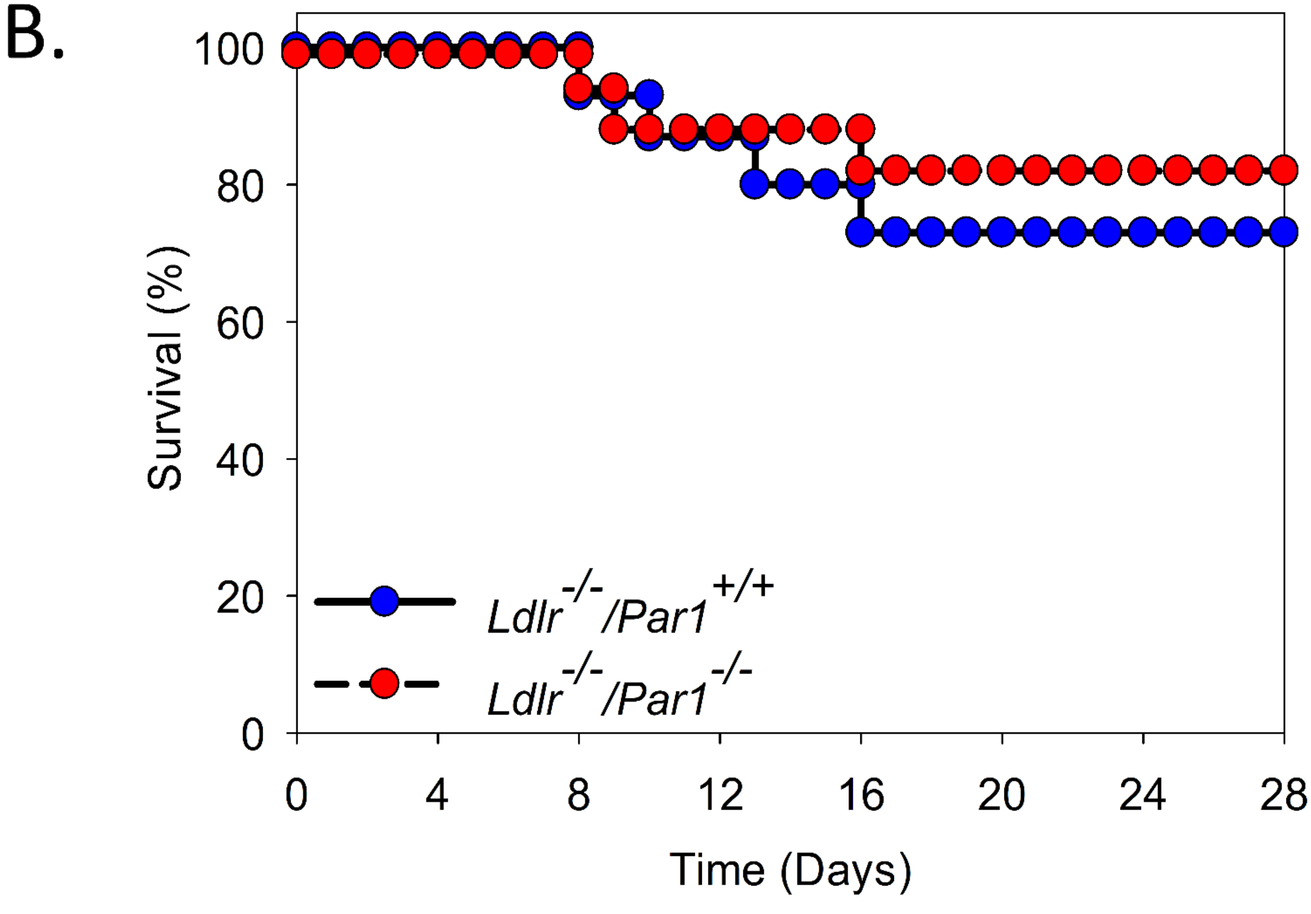

**Supplemental Figure 5**

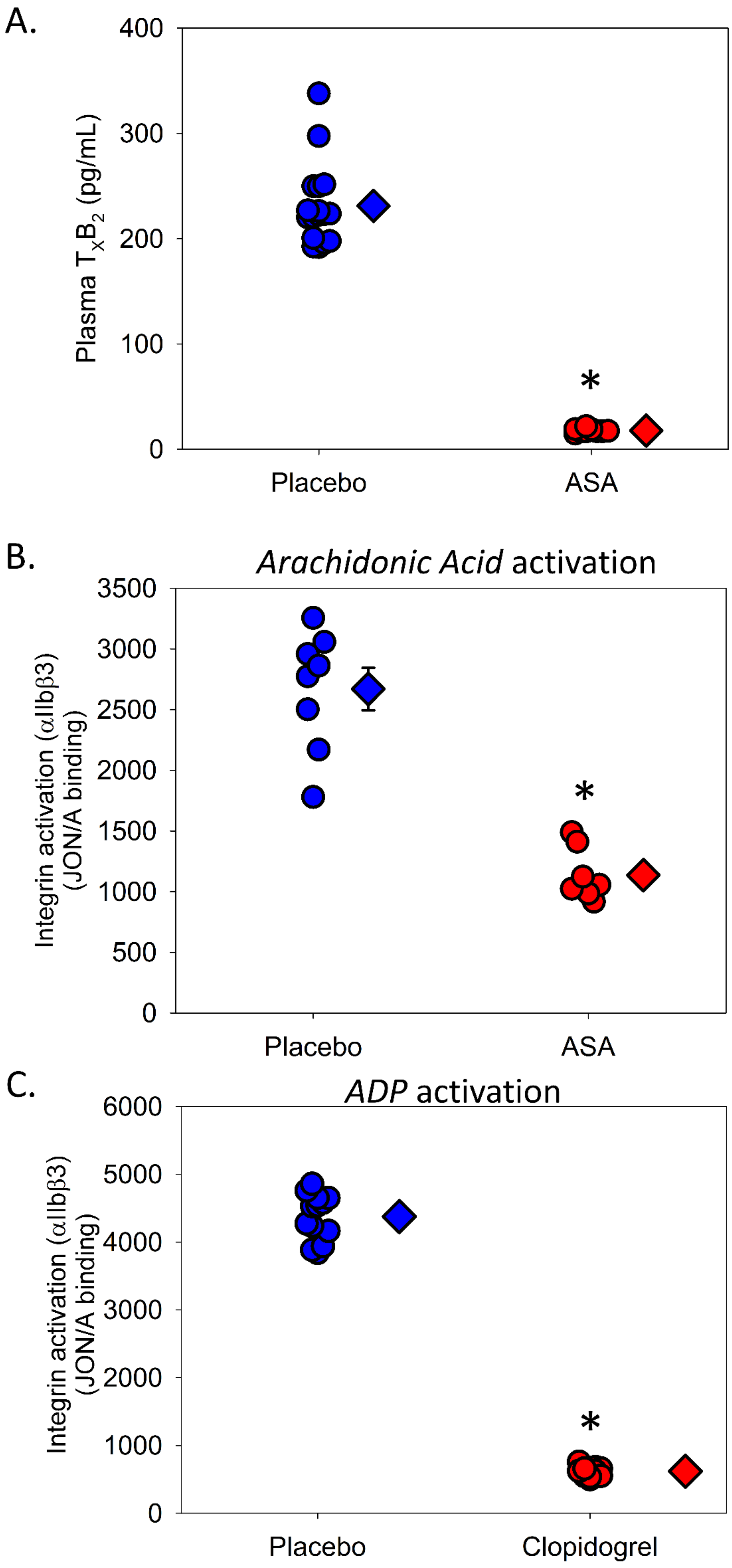

**Supplemental Figure 6**

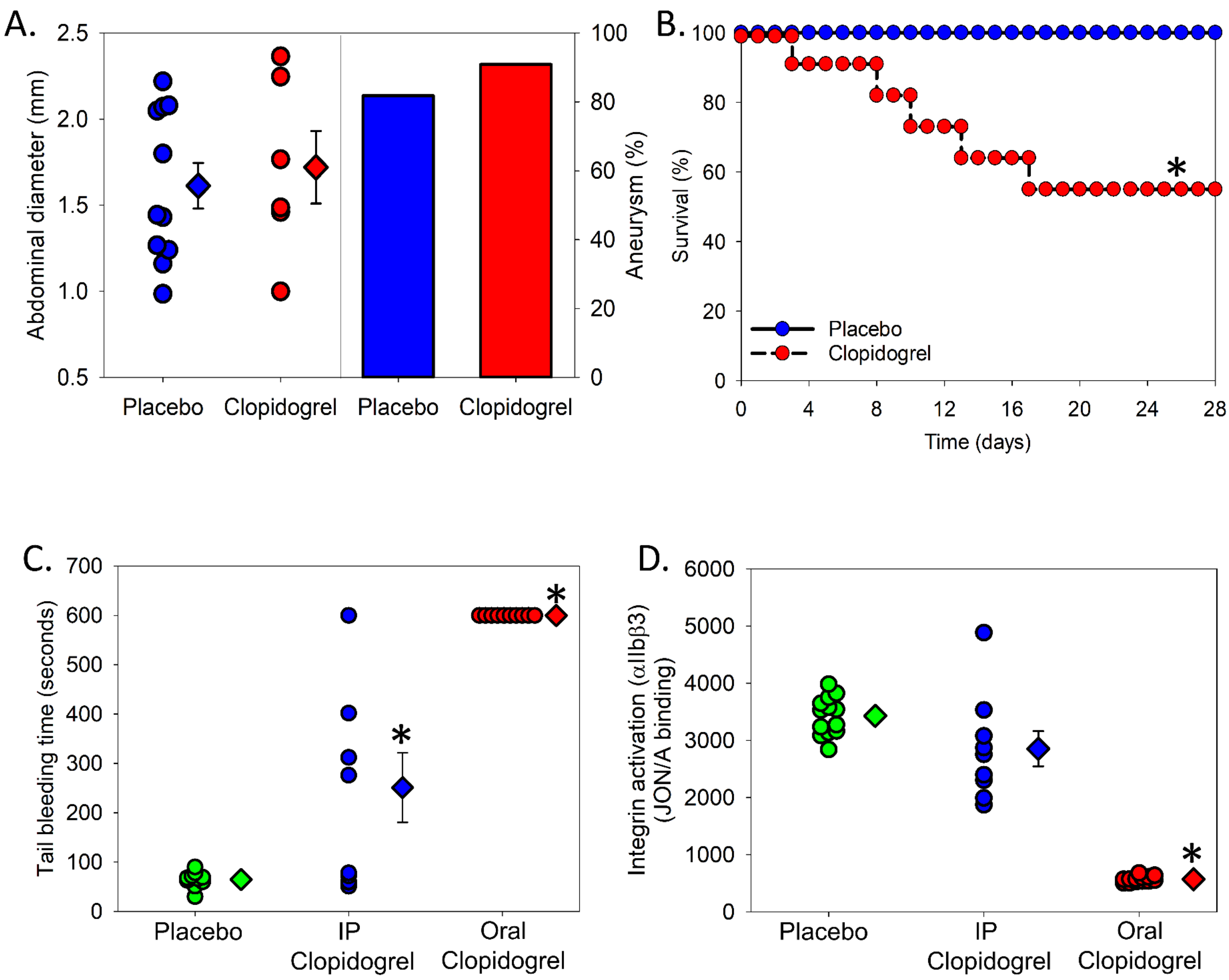
